# Brain glucodynamic variability is an essential feature of the metabolism-cognition relationship

**DOI:** 10.1101/2025.04.30.651418

**Authors:** Hamish A. Deery, Emma Liang, Chris Moran, Gary F. Egan, Sharna D. Jamadar

## Abstract

Variability ensures that complex biological systems, including the brain, are capable of responding to changing environmental demands. While the importance of neural variability in electrophysiological and haemodynamic aspects of brain activity is beginning to be understood, little is known about how variability in molecular activity influences brain function. Here we examine how temporal variability in *glucodynamics*, or time-varying glucose use, is related to cognition in 35 younger and 43 older adults. Stationary metabolic rates of glucose were not directly associated with cognition. Rather, higher glucodynamic variability, and its coherence into metabolic networks, was associated with better cognitive performance. Lower glucodynamic variability in ageing was associated with altered metabolic network efficiency and reduced cognitive performance. Our results demonstrate for the first time that variability in cerebral glucose metabolism is biologically and functionally relevant to cognition and the network architecture of the brain. Cognition is influenced by time-varying glucose metabolism and its coherent fluctuations in metabolic networks. A loss of glucodynamics in ageing reduces the efficiency of the metabolic connectome and contributes to reduced cognitive performance. The study of glucodynamics significantly advances our understanding of metabolic brain changes in health, ageing and disease.

**Significance statement:** The human brain has a high energy requirement and must be able to dynamically adjust its energy supply as needed. However, the dynamic nature of brain glucose use is not well understood. In this work, we identify the variability in glucodynamics - the moment-to-moment fluctuations in glucose use - as an important biological feature supporting cognition and mediating age-related changes in brain function. We report that higher glucodynamic variability is associated with higher metabolic network efficiency and enhanced cognitive performance. Older adults exhibit reduced glucodynamic variability, reduced network efficiency and impaired cognition. Our results demonstrate that glucodynamics are important to our understanding of energy-dependent cognitive processes, the efficiency of metabolic brain networks, and the energetic basis of cognitive ageing.

## Introduction

The human brain uses glucose as its primary metabolic substrate to support its high energetic demands [1-5]. Neuronal activity depends on a continuous glucose supply to support energy-intensive processes, such as the generation of action potentials, neurotransmitter release, and synaptic signalling [6]. Glucose availability also shapes large-scale brain network architecture and cognitive function [1, 7]. Given the brain’s limited capacity for glucose storage, it must dynamically regulate energy availability to sustain the metabolic demands associated with neuronal activity and adaptive cognitive processing. However, the mechanisms by which glucose metabolism is modulated in real time to meet the energetic requirements of cognition remain poorly understood [8].

Variability is an essential feature of complex biological systems [9]. Variability ensures that systems are flexible, adaptive and capable of responding to changing internal and external demands. Neuronal variability, especially in synaptic strength and firing patterns [10-12], is necessary for the brain’s ability to learn from experience and adapt to new situations [13-15]. Coherent fluctuations in neuronal activity give rise to connectivity between regions in large-scale networks [16]; this is thought to underlie information transfer in the brain [17, 18]. Intrinsically-generated variability in neural activity allows different regions of the brain to become more or less connected depending on ongoing processing needs [19]. Dysfunction in dynamic neural activity and connectivity patterns can lead to cognitive deficits and is a major contributor to the development of neurological and psychiatric conditions [20-22].

Variability in neuronal fluctuations that underpin network connectivity are thought to represent the information processing capacity of the system, with higher variability reflecting greater capacity for information integration across the network [23]. At the macroscale, variability in brain function has been measured using the time-varying blood oxygenated level dependent (BOLD) functional magnetic resonance imaging (BOLD-fMRI) signal (timeseries). Regional BOLD variability within individuals maps to known functional network topology and greater BOLD variability is associated with better cognitive performance [24, 25]. Conversely, increasing age has been associated with a loss of BOLD signal variability and worse cognitive performance [15, 24-27]. Similar age and cognitive effects have been associated with the variability of micro- and macro-scale electrophysiology [21, 28-30].

A limitation of the BOLD signal is that it is an indirect measure of neuronal activity and is influenced by multiple components of metabolism [1, 31], confounding the interpretation of the metabolic basis of the signal [32, 33]. In contrast, glucose metabolism measured with (18F)-fluorodeoxyglucose positron emission tomography (FDG PET) provides a more direct measure of glucose metabolism at the post synaptic neuron [1]. FDG PET has traditionally been used to measure brain function under the assumption of a steady state [34], and thereby disregards temporal changes of glucose metabolism within individuals. Recent methodological advances in time-resolved PET [34, 35], including functional FDG PET (fPET), allow for dynamic glucose metabolism to be estimated for individuals and for time-varying metabolism to be studied [36-40], which we call *glucodynamics* [1]. Regional glucodynamic signals show coherence, forming a ‘metabolic’ functional connectome of the brain ([36, 38, 41-43], see also [44-46]).

However, the temporal variability of glucose metabolism within individuals is yet to be studied. A knowledge of variability in cerebral glucose metabolism is particularly important to understand how the brain adapts to changing conditions and how energy supply and use influences cognition and behaviour [1]. The connectivity strength of the brain’s metabolic connectome reduces with age, becoming reconfigured to be more integrated and reliant on posterior hubs. This pattern comes at a high metabolic cost and is associated with worse cognitive performance [43]. Reduced moment-to-moment glucose availability and/or variability in glucose metabolism may explain some of the loss of metabolic network coherence and efficiency seen in ageing. The loss of variability in glucose metabolism likely limits the brain’s ability to switch between network configurations due to reductions in coherent neural activity. This then leads to less efficient information transfer between networks [27, 47, 48].

Here we study changes to the variability of glucodynamics in normative ageing, measured as the standard deviation of the time-varying fPET timeseries for individuals. We examine the associations of variability within individuals with a ‘stationary’^1^ measure of glucose (i.e., CMR_GLC_ indexed across the entire scan), metabolic network efficiency and cognitive performance. Based on evidence from EEG [29, 49] and fMRI [15, 24, 25] studies, we hypothesise that the variability of the fPET signal will be lower in older than younger adults. We also hypothesise that lower fPET variability will be associated with lower local and global metabolic network efficiency. Finally, we hypothesise that fPET signal variability mediates the association between CMR_GLC_, network efficiency and cognition.

## Results

The Monash MetConn simultaneous PET/MR dataset [32, 43] of N=78 healthy younger and older adults was used in this study (Table 1; Supplement 2.1). Stationary metabolic rates of glucose (CMR_GLC_) across the 90 min scan were calculated using Patlak modelling. fPET data was reconstructed with 16s frame duration and glucodynamic variability calculated using the standard deviation of the fPET timeseries. Figure 1 shows an example fPET timeseries and fPET_SD_ from one region for a single participant. The metabolic connectome was constructed from the fPET timeseries using Pearson correlations among 100 functional regions, with network topology characterised using global and local efficiency. Finally, cognitive data to assess proactive and reactive control, episodic memory and processing speed was reduced in dimensionality using Principal Component Analysis (PCA), to yield a single component explaining 58% of the variance (see Supplement 2.2). Older adults showed worse performance on the cognition PC (Figure 2b.v).

**Figure 1.**
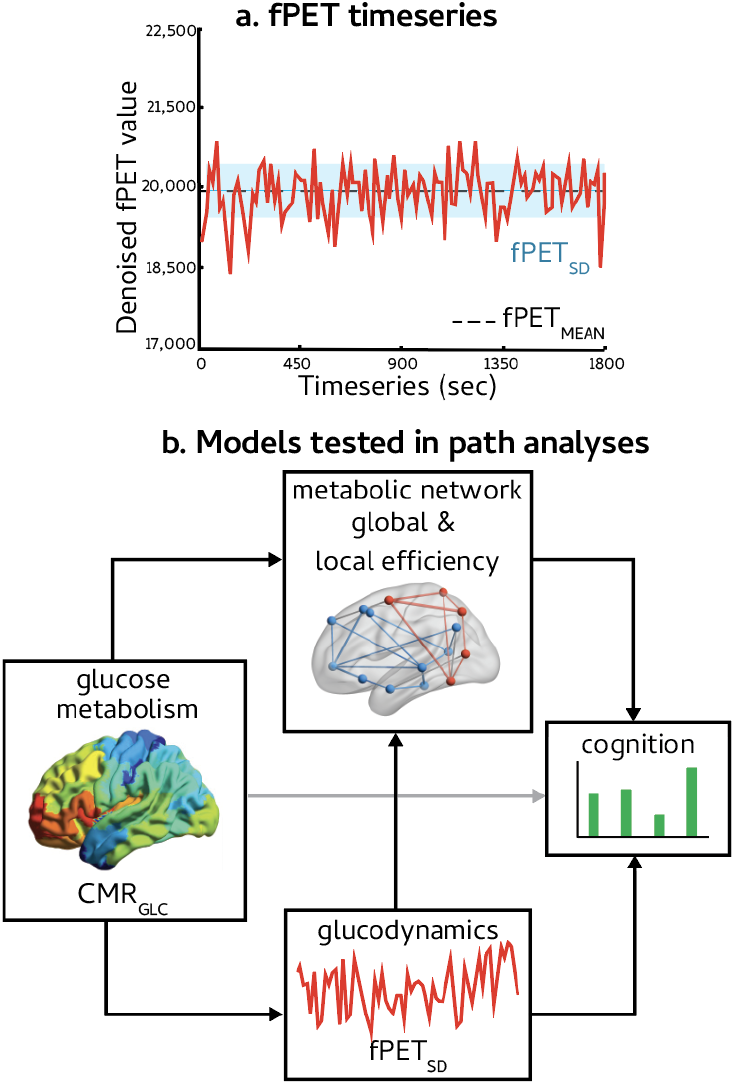
Illustration of fPET timeseries and models tested in the path analyses. The fPET timeseries (A) for one subject in the left visual striatal cortex, with blue shading illustrating the standard deviation of the timeseries, i.e., fPET_SD_, indexing ‘glucodynamics’ across the scan. Separate path analyses (B) were tested for the whole brain and the dorsal attention, salience ventral attention, control and default networks. The path analyses included direct effects of fPET_SD_ on cognition and indirect effects of CMR_GLC_ and fPET_SD_ on cognition via local and global network efficiency. CMR_GLC_ was indexed as a ‘static’ measure of glucose metabolism across the entire scan. Models were also tested in which there was a direct path from CMR_GLC_ to cognition (grey arrow).

**Figure 2.**
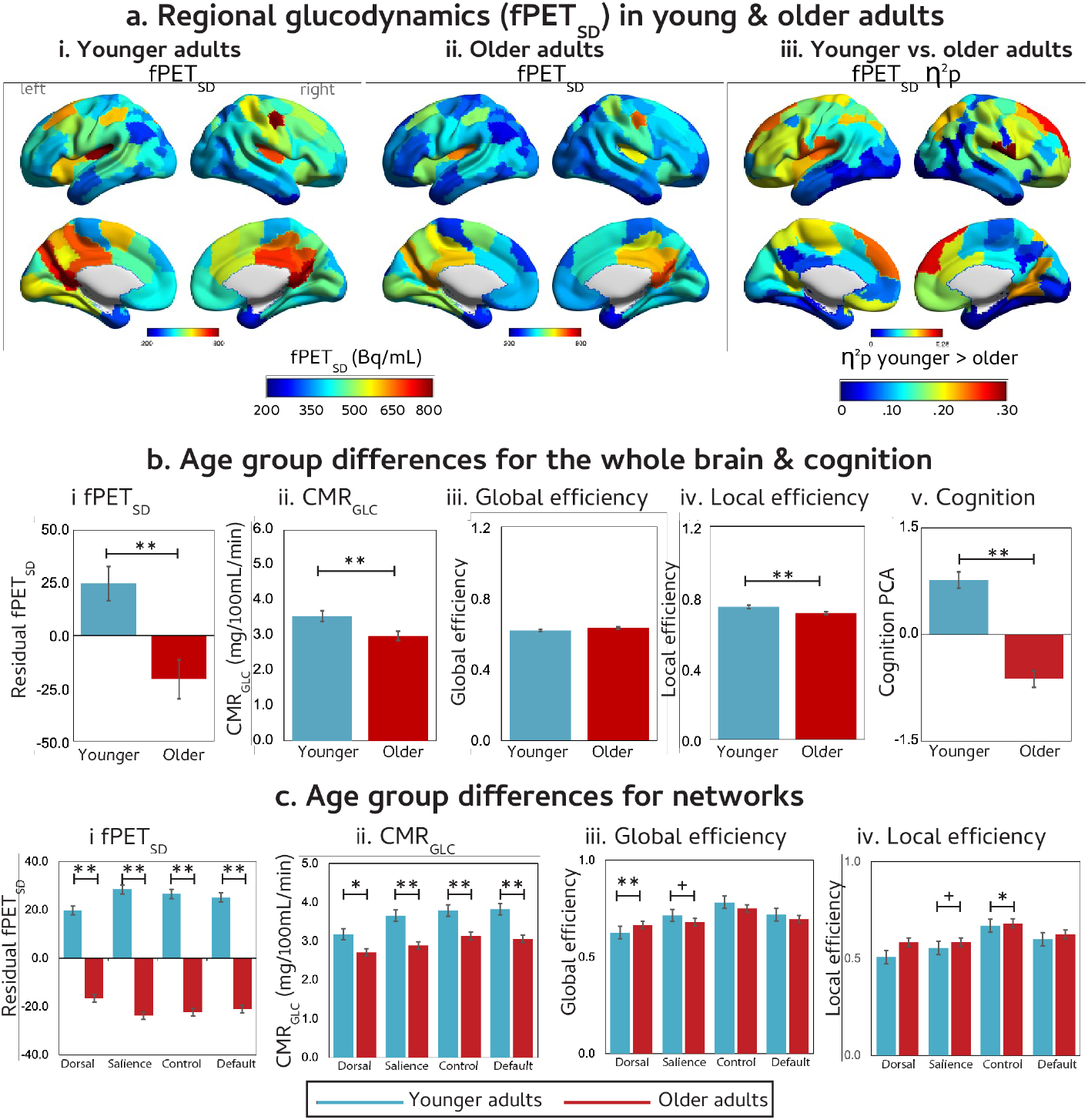
Age group differences in fPET_SD_, brain network measures and cognition. (A) Mean regional fPET_SD_ for younger (i) and older (ii) adults; Effect sizes (η^2^_p_) of higher regional fPET_SD_ in younger versus older adults (iii). There was no region in which older adults had higher fPET_SD_ than younger adults. Head motion was included as a covariate. (B) Age group means (and standard errors bars) and t-test of age group differences in whole brain fPET_SD_ (i), CMR_GLC_ (ii), local and global efficiency (iii and iv) and cognition (v). (C) Age group means (and standard errors bars) and t-test of age group differences in network fPET_SD_ (i) CMR_GLC_ (ii), and local and global efficiency (iii and iv). fPET_SD_ and CMR_GLC_ were indexed as ‘glucodynamic’ and ‘static’ measure of glucose use and metabolism across the entire scan. p-FDR < 0.05; **p-FDR < 0.01; ^+^p< 0.05 before FDR-correction. DA = Dorsal Attention’ SA = salience ventral attention; CON = control; and DEF = default network. Age group means and effect sizes are available in Supplement 2.4, 2.7 and 2.8. Figures were produced using Brain Net Viewer (http://www.nitrc.org/projects/bnv/) (Xia et al., 2013).

Examination of the fPET timeseries indicated that older adults show lower glucodynamic variability (fPET_SD_) and reduced efficiency of the metabolic connectome relative to younger adults (Figure 2a and 2b.i). Older age was associated with lower fPET_SD_ in 74 regions. There was no region in which older adults had higher fPET_SD_ than younger adults. Across the whole brain, older adults also had lower CMR_GLC_ compared to younger adults (Figure 2b.ii), replicating previously reported results [50]. Older adults had lower local efficiency of the metabolic connectome than younger adults (Figure 2b.iv). Although the older adults tended to have higher global efficiency in their metabolic connectome than younger adults, the difference was not statistically significant (Figure 2b.iii).

We examined whether these whole brain effects were maintained at the functional sub-network level. We focused specifically on the associative networks (dorsal attention, salience ventral attention, control and default mode networks) as our previous report demonstrated that the fPET signal in these networks was most strongly associated with cognitive outcomes [32, 43]. Older adults had lower fPET_SD_ and CMR_GLC_ than younger adults in all networks (Figure 2c.i and 2c.ii). Across each of the networks, the largest between-age groups fPET_SD_ differences were in the prefrontal and parietal regions. Older adults had lower local efficiency in the control metabolic network than younger adults (Figure 2C.iv); whereas global efficiency in the dorsal attention network was greater in older adults (Figure 2C.iii). This pattern of results is consistent with the reduced segregation (or modularity) and increased integration in metabolic [32, 43] and functional fMRI connectomes [51] seen in older age.

To understand how glucodynamic variability influences cognition, we used path analysis. We investigated the relative contributions of glucodynamic variability (fPET_SD_), stationary (entire scan) metabolic rates of glucose (CMR_GLC_) and the efficiency of metabolic network to cognition. The model is shown in Figure 1b. Strikingly, when the direct effect of CMR_GLC_ on cognition (grey line in Figure 1b) was included in the model, the model fit was worse relative to models with an indirect effect only of CMR_GLC_ on cognition via glucodynamics and network efficiency (see Supplement 2.5 for goodness-of-fit). This was the case for the whole brain and each of the network models. Moreover, in those models, the direct path from stationary glucose metabolism to cognition was weak (betas <.12) and non-significant (p>.280). We therefore focused our analyses on models in which the effect of CMR_GLC_ on cognition was indirect via glucodynamics and network efficiency.

For the whole brain, greater availability of glucose (higher CMR_GLC_) was associated with greater glucodynamic variability (higher fPET_SD_) and better cognitive performance (red paths in Figure 3a). Higher CMR_GLC_ was also indirectly associated with better cognitive performance via network efficiency (blue paths in Figure 3a). Specifically, higher CMR_GLC_ was associated with greater fPET_SD_, which was associated with greater local (Figure 3ai) and lower global (Figure 3aii) efficiency and better cognitive performance (Figure 3a blue lines).

**Figure 3.**
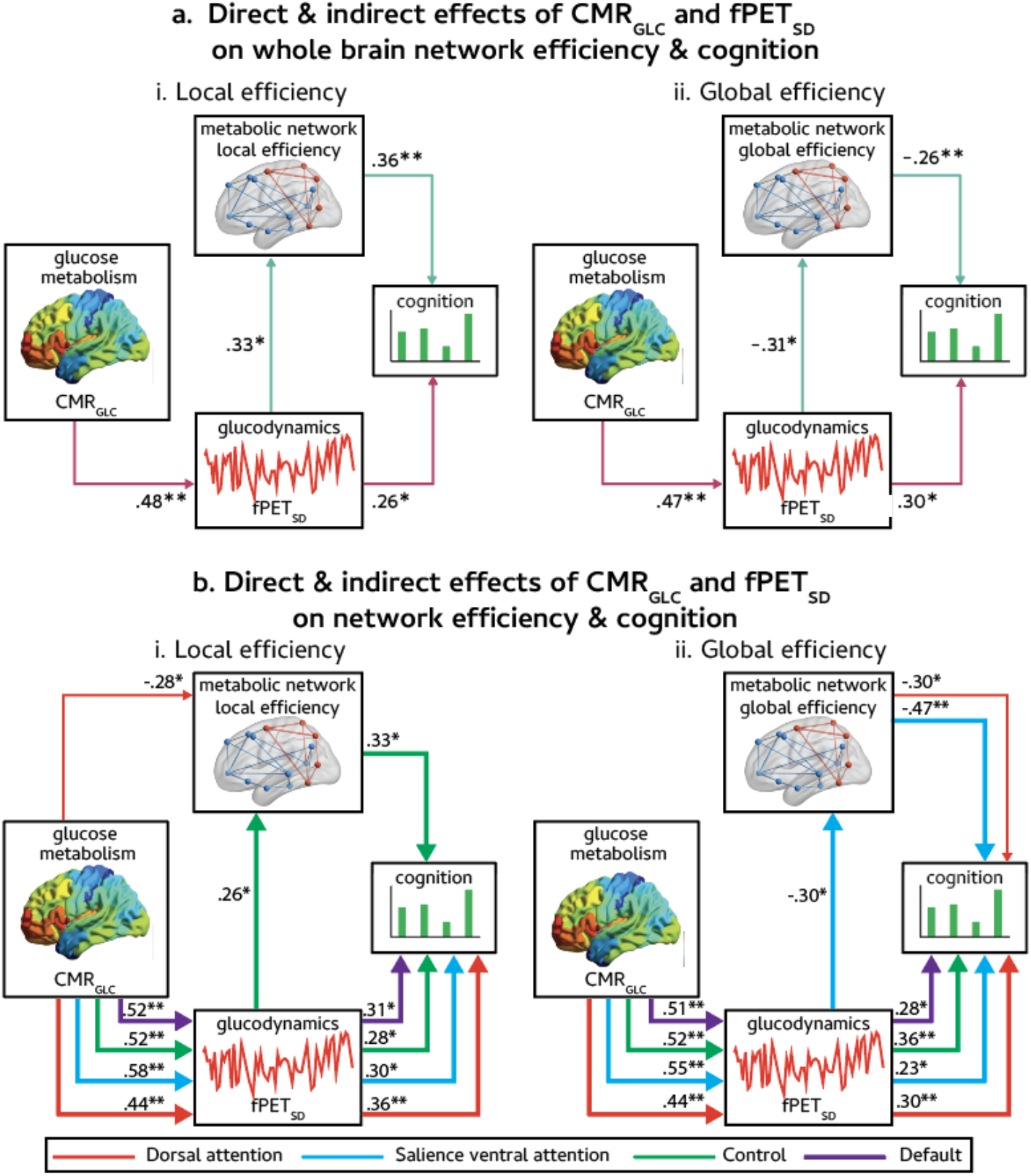
Path analyses testing direct and indirect effects of CMR_GLC_ and fPET_SD_ on network efficiency and cognition. Path analyses of whole brain (A) and network-level (B) measures including local (Ai and Bi) and global efficiency (Aii and Bii). Numbers on the arrows are standardised beta weights. In each model, only the statistically significant paths are shown. Thicker lines illustrate where there is a significant association between CMR_GLC_ and cognition, including via fPET_SD_ and/or network efficiency. *p-FDR < 0.05; **p-FDR < 0.01; ^+^p< 0.05 before FDR-correction. The t-test of age group differences in the measures used in the path analyses are available in Supplement 2.4, Full details of all paths analyses, including the chi-square values for goodness-of-fit, regression weights and their significance levels are in Supplement 2.6.

Effects of glucodynamics on cognition were seen at the network level, albeit with some different effects on cognition via network efficiency. All four networks showed a direct association between CMR_GLC_, fPET_SD_ and cognition, indicating the importance of glucodynamic variability on cognition. In all four networks, greater availability of glucose (higher CMR_GLC_) was associated with greater glucodynamic variability (higher fPET_SD_) and better cognition (purple, green, blue and red lines in the lower half of Figure 3b). In the control network, higher CMR_GLC_ was associated with higher fPET_SD_, higher local efficiency and better cognitive performance (Figure 3B.i). In the salience ventral attention network, higher CMR_GLC_ was associated with higher fPET_SD_, *reduced* network global efficiency and *better* cognitive performance. Finally, the dorsal attention network showed two significant paths that did not link stationary glucose metabolism to cognition. Greater availability of glucose in the dorsal attention network was associated with reduced local efficiency of that network (Figure 3b.i red arrow in top left quadrant). This was the only path between stationary glucose metabolism and network efficiency in any model. Reduced global efficiency of the dorsal attention network was associated with better cognition (Figure 3b.ii red arrow top right quadrant), however this was unrelated to CMR_GLC_ or fPET_SD_ in that network.

## Discussion

This study demonstrates the crucial role that glucodynamic activity plays in cognition and cognitive ageing. Time-varying glucose use (‘glucodynamics’) and its coherence into metabolic networks directly influences cognition and influences the link between stationary glucose metabolism and cognition. Our results indicate that variability in cerebral glucose metabolism is biologically and functionally relevant to cognition and the network architecture of the brain.

The majority of the energy budget of the brain is dedicated to maintaining the spontaneous activity of the resting state [1, 3, 52]. Variability in this spontaneous activity reflects both the cytoarchitecture of the brain region, its current association with a functional subsystem (which may change over time [53, 54]), as well as its centrality and hubness, or importance in the network [1]. Variability in spontaneous activity can be considered an index of the brain’s ‘dynamic range’ of responses, with an optimal dynamic range supporting flexible and adaptive behavioural responses [10]. Here, a larger energy budget (CMR_GLC_) was associated with greater dynamic range (fPET_SD_) and better cognition, at both the whole brain and brain network level. This pattern was robust and was evident across all sub-networks we studied. Our results are consistent with the argument that neural variability is fundamentally important for information propagation throughout the network [55].

The direct path from glucodynamics to cognition suggests that variability in brain metabolism is directly linked to cognitive performance. However, at the whole brain level, there was also an indirect path linking glucodynamics to cognition through the network efficiency of the metabolic connectome. This indicates that glucodynamic variability is not the only contributor to cognition, but the coherence of glucodynamic fluctuations between regions that form networks is important as well. Network-level analyses indicated that the prediction of cognition from coherent network dynamics was selective for two networks: namely, the control network and the salience ventral attention network. The cognitive battery used here specifically targeted proactive and reactive cognitive control, processing speed and verbal episodic memory – tasks that rely substantially on the control and salience networks. As such the finding that the efficiency of the control and salience ventral attention networks is important for cognition may be somewhat unsurprising. It is notable, however, that foundational work on the regulation of glucose availability on cognition indicates that these networks, which support working memory and inhibitory control, are particularly sensitive to the availability of glucose in the system [56]. Increasing the availability of glucose profoundly improves working memory and inhibitory control, particularly in ageing when glucose availability decreases ([43]; also see [56] for a review). Our results complement and significantly extend earlier work, indicating that not only glucose availability but also the dynamic and coherent use of glucose shapes control and inhibitory cognitive processes.

Network efficiency reflects the brain’s ability to transfer and process information across different brain regions. Interestingly, higher fPET variability and better cognitive outcomes were associated with *higher* local efficiency and *lower* global efficiency. Together, the measures of local and global efficiency provide insight into the balance of network integration and segregation of the metabolic network. The human functional brain network is organised into modules or sub-networks of functionally connected regions [57], with local efficiency indexing the efficiency of communication within a sub-network (local integration) and global efficiency indexing the efficiency of information transfer across the entire network (global integration). Recent evidence indicates that the metabolic network is similarly organised [32, 43]. A healthy balance of segregation and integration of functional sub-networks supports flexible and adaptive responses, and disruption of this balance leads to deficient information transfer across the network, reduced capacity to deal with complexity, and a sub-optimal use of neural energy [43[Vazquez-Rodriguez, 2017 #57, 57]. The positive association between glucose availability (CMR_GLC_), glucodynamics (fPET_SD_), local efficiency and cognition may reflect the brain prioritising metabolic resources for local information processing among regions that are topologically close, which is less energetically demanding than information transfer via long-range connections [58]. This topology minimises the metabolic cost of local information processing, allowing for fast and efficient signal transmission. On the other hand, the negative association between glucose availability, glucodynamics, and global efficiency may reflect an increased cost of supporting long-range connections across a more globally integrated network [1, 57, 59].

Higher stationary glucose metabolism was indirectly associated with better cognition via higher network glucodynamics and via network efficiency. However, these metabolism-cognition relationships cannot be adequately explained by the time-invariant measure of glucose metabolism. In fact, a direct effect of CMR_GLC_ on cognition resulted in a poorer model fit and in weak and non-significant associations of CMR_GLC_ and cognition. These results indicate that glucodynamics and their regional coherence indexed by network efficiency provide a more complete understanding of the metabolism-cognition relationship than does CMR_GLC_ as a single measure of metabolism. These results also suggest that studies of cerebral metabolism need to measure glucose within individuals and index variability of the timeseries to fully understand the role of glucose in the brain.

### Loss of glucodynamic variability in healthy ageing

Older adults show lower stationary glucose metabolism across the cortex than younger adults. This is an established finding (see [50] for a review) and indicates that older adults have a reduced budget of glucose to support neural functions compared to younger adults. Importantly, we also found that older adults display reduced glucodynamic variability and poorer cognitive performance, indicating that ageing is also associated with a pervasive loss of the dynamic range of glucose metabolism. A loss of glucodynamic range is compatible with previous results indicating that a loss of electrophysiological and haemodynamic variability occurs in ageing and is associated with poorer cognitive outcomes [10, 15, 60]. A loss of brain signal variability in ageing has been hypothesised to reduce the capacity of the brain to shift between segregated and integrated functional brain configurations [26]. This is consistent with our finding that older adults have lower metabolic network local efficiency and higher global efficiency compared to younger adults and worse cognitive performance.

We previously reported [43] that the metabolic connectomes of older adults are distinguished by reduced metabolic connectivity in the frontal, motor, parietal and medial cortices. Here we found older age was associated with particularly lower glucodynamics across the prefrontal cortex, parietal lobule and several regions of the somatomotor network. Taken together, these results suggest that changes in metabolic connectomes of older adults, particularly connections in the frontal regions, are due to reduced glucose availability and variability. The mechanisms underpinning these changes in ageing are likely to act at multiple spatial and temporal scales. For example, there is a loss of efficiency of regional metabolism in ageing, even after adjusting for atrophy in older adults [50, 51], and a higher glucose cost to maintain large-scale network connectivity [43]. At a microscale, the ratio between synaptic excitation and inhibition influences the efficiency of energy consumption and information transmission of neural networks [61] and is attenuated in ageing [61]. The FDG-PET signal primarily reflects post-synaptic activity [62]. Postsynaptic variability may reflect variability in the signal received from the presynaptic spike train [63]. Presynaptic spike trains encode information through the number of spikes, the spike rate, and the inter-spike interval [10], and abnormalities in spike dynamics and oscillatory activity occur in ageing [64].

The idea that a loss of glucodynamics in ageing may lead to a reconfiguration of the metabolic connectome is consistent with theories of cognitive ageing that have focused on the overall efficiency in the recruitment and use of neuronal resources [65]. The *scaffolding theory of ageing and cognition* [66] holds that the recruitment of additional or different neural resources via network reorganisation provides the foundation to maintain cognitive function in the face of age-related structural and functional decline. In this model, a more integrated system in older age can be considered compensation for a change in the underlying structural and functional “scaffolding” of the brain [67]. Our results suggest that a loss of glucose availability and glucodynamics also contribute to changes in the brain’s network organisation in ageing.

### Glucodynamic variability is an important dimension of brain variability

The idea that biological variability is necessary for optimal brain function is central to models of *coordination dynamics* [68, 69], which proposes that the brain maintains a *meta-stable* state, where cooperative (integrative) and independent (segregative) processes co-exist [68-70]. Rather than existing in a state where groups of brain regions react to an event and then return to some homeostatic equilibrium ready to react to the next event; brain regions in a meta-stable state are continuously engaged in multiple interactions, with changes occurring at any place and point in time, and reverberations through the network that affect other regions [54, 68]. Consequently, some brain regions work together and coordinate their activity, whereas other brain regions remain relatively autonomous and complete specialised functions [70]. A healthy dynamic range of functional brain activity supports this dynamic meta-stable state. Our results highlight that biological variability occurs not only on electrophysiological and haemodynamic scales of functional connectivity, but also on metabolic scales – consistent with the idea that the brain’s functional interactions are multiple, with information transferred across various biological dimensions (electrophysiological, molecular, haemodynamic) between nodes of the network [53].

It is important to note that while the standard deviation of the fPET timeseries does provide information about the dynamic nature of glucose utilisation [24, 25], there are limitations in the richness of the dynamic information. To demonstrate meta-stability, and not related states such as multi-stability, formal modelling based on dynamical systems theory is required [70.]. Further, we have noted that while traditional scan-wise measures of CMR_GLC_ assume weak sense stationarity (see Footnote 1), or a stable and non-changing rate of glucose metabolism across the scan, there is an element of this assumption in the timeseries analysis applied here [71]. We indexed the time-varying glucose utilisation across the scan, and used it to estimate a single scan-wise metabolic connectome for each individual, as well as the variability of the signal across the scan. As with the vast majority of functional connectivity studies using fMRI, the analysis assumes that the connectivity and the variability is unchanging across the scan [71, 72], differing from fully dynamic approaches, such as time varying connectivity and sliding window approaches [73]. The capacity to delineate a subject-level metabolic connectome and capture glucodynamics are recent technical advances [35, 38, 39, 43, 74] and so there is still much to be learned about time-varying neural glucose utilisation. An important future direction of this research is to expand the study of glucodynamics to capture the temporal pattern of glucose use and its coherence into network states. This will enable examination of phenomena such as microstates, dwell states, and quasi-periodic patterns: periods of stability in the brain’s ongoing dynamic activity.

Our study has limitations. The path analyses suggest a causal chain from CMR_GLC_ to cognition via glucodynamics and network efficiency. Techniques like metabolic connectivity mapping and dynamic causal modelling can validate the temporal direction of our models by assessing how metabolism in a target region is driven by incoming signals from other regions [46]. Longitudinal research can assess whether the timing of changes in glucodynamics correspond to changes in network alterations and cognition. In the current study, glucodynamics were measured at a temporal scale of 16 seconds. This represents a step-change in temporal resolution by comparison to other PET acquisitions (which are usually acquired at steady state over 10-30mins), but is slow by comparison to other methodologies. The biological consequences of neural variability and meta-stability are believed to be present at all scales of space and time [53, 68, 69]. At longer timescales, function is likely to reflect the underlying structural constraints of the network [53, 75], which are themselves constrained by the physical space of the brain and the energetic constraints of the system [3, 59, 76]. Biologically, the extraction of glucose from the bloodstream to support neuronal function occurs at a fast timescale, with around one molecule of glucose extracted at the timescale of a synaptic event [6]. It is possible that future technical advancement in measuring glucodynamics at even higher temporal resolution will uncover new insight into the functional meaning of glucodynamic variability and its role in ageing and disease.

The human brain is a highly metabolically active organ. It relies upon a reliable and *scalable* (i.e., able to be up- and down-regulated on demand) supply of glucose to support its function. How brain metabolism is regulated on demand to meet the energetic requirements of cognition remains an open question [8]. Here we show that variability in the moment-to-moment metabolism of glucose is biologically and functionally important to cognition and in explaining metabolic and brain network changes in ageing. Glucodynamics are a fundamental feature of the human brain’s functional network organisation. The link between glucose and cognition is attributable to glucodynamics and its coherence into metabolic networks. The integration and efficiency of the metabolic network is realised via the synchronised glucodynamics of regions and is altered in ageing. Variability in glucodynamics also directly predicts cognitive performance and indirectly predicts cognition via network efficiency. As such, glucodynamics appear to index the information processing capacity of functional brain networks. Studying glucodynamics offers promise as a means to increase our understanding of metabolic brain changes in health, ageing and disease states.

## Methods

Full details of the Methods are provided in Supplement 1.

### Participants

The final sample included 78 individuals, 35 younger (mean 27.8; SD 6.4; range 20-42 years) and 43 older (mean 75.4; SD 5.7; range 66-86 years) adults (see Supplement 2.1).

### Data Acquisition

#### Cognitive Tests

Participants completed an online demographic and lifestyle questionnaire and a cognitive test battery. The cognitive domains and tests were chosen because of their well-established validity in predicting cognitive changes in ageing (see for [77, 78] review). The following measures were used: Hopkins Verbal Learning Test (HVLT), task-switching, stop-signal, and digit symbol substitution. Principal components analysis was used to reduce dimensionality in the cognitive data, yielding one significant PC explaining 58% of the variance.

#### PET Data Acquisition

Participants underwent a 90-minute simultaneous MR-PET scan in a Siemens (Erlangen) Biograph 3-Tesla molecular MR scanner. At the start of the scan, half of the 260 MBq FDG tracer was administered as a bolus to provide a strong PET signal. The remaining 130 MBq of the tracer dose was infused over 50 minutes at a rate of 36ml/hour. This combined bolus plus constant infusion protocol provides a good balance between a rapid increase in signal-to-noise ratio at the beginning of the scan, and maintenance of signal-to-noise ratio over the length of the scan [42].

The scan sequence commenced with non-functional T1 and T2 MRI scans in the first 12 minutes to image the anatomical grey and white matter structures, respectively. The T1 3DMPRAGE scan parameters were: TA = 3.49 min, TR = 1,640ms, TE = 234ms, flip angle = 8°, field of view = 256 × 256 mm^2^, voxel size = 1.0 × 1.0 × 1.0 mm^3^, 176 slices, sagittal acquisition. The T2 FLAIR parameters were: TA = 5.52 min, TR = 5,000ms, TE = 396ms, field of view = 250 × 250 mm^2^, voxel size = .5 × .5 × 1 mm^3^, 160 slices). Thirteen minutes into the scan, list-mode PET (voxel size = 1.39 × 1.39 × 5.0mm^3^) and T2* EPI BOLD-fMRI (TA = 40 minutes; TR = 1,000ms, TE = 39ms, FOV = 210 mm2, 2.4 × 2.4 × 2.4 mm^3^ voxels, 64 slices, ascending axial acquisition) sequences were initiated. A 40-minute resting-state scan was undertaken while participants watched a movie of a drone flying over the Hawaii Islands.

### MRI & PET Pre-Processing

For the T1 & T2 images, the brain was extracted in Freesurfer and registered to MNI152 space. The quality of the pial and white matter surface was manually checked and corrected if needed.

Participants’ list-mode PET data was binned into 344 3D sinogram frames at 16s intervals. Attenuation was corrected via the pseudo-CT method for hybrid PET-MR scanners [79]. 3D volumes were constructed from the sinogram frames using the Ordinary Poisson-Ordered Subset Expectation Maximization algorithm (3 iterations, 21 subsets) with point spread function correction. The 4D PET volumes were motion corrected [80], using the mean PET image to mask the data. PET images were corrected for partial volume effects using the modified Müller-Gartner method [81]. Regions of interest were generated for the 100 regions of the Schaefer Atlas [82].

### Derivation of metabolic rate of glucose, glucodynamics, and the metabolic connectome

Regional CMR_GLC_ was indexed as a ‘stationary’ measure of glucose metabolism across the entire scan. Calculations of regional CMR_GLC_ were undertaken using the FDG time activity curves for the Schaefer atlas regions [83] using Patlak modelling.

fPET_SD_ was calculated from the individual participant’s 100 regional timeseries. Data from 20 to 50 minutes of the scan were used to reduce the impact of the initial FDG uptake ramp and decrease post cessation of the constant infusion [41]. fPET_SD_ measures the dispersion or spread of the signal around the mean, reflecting variability across the timeseries (see Figure 1A for illustration).

The topology of the metabolic connectome was described by the global and local efficiency graph metrics calculated at the top 30% of network edges based on relative edge strength. Local efficiency at each node is the average of shortest inverse-distances between the node and its neighbouring node in the local sub-graph. Global efficiency at each node is the average of the shortest inverse-distances between the node and all other nodes in the entire graph. Local and global efficiency across the whole graph are measures of local and global integration, respectively [17]. Global and local efficiency were calculated for the whole brain graph, as well as for the networks of the sub-graph comprising the dorsal attention, salience ventral attention, control and default networks.

### Data Analysis

To test our hypothesis that older age is associated with lower variability of the fPET timeseries, a series of general linear models (GLMs) was run in which regional fPET_SD_ was the dependent variable and head motion was a covariate. Partial eta squared (η^2^_p_) was used to quantify the age effect sizes in the GLMs. Each main effect was FDR-corrected at p < .05 across the 100 regions.

We used path analyses to test our hypotheses that that lower fPET_SD_ will be associated with lower network efficiency and that fPET_SD_ mediates the associated between ‘stationary’ glucose metabolism, network efficiency and cognition. A separate path analysis was undertaken at two spatial scales: 1) the whole brain; and 2) each of the four networks separately: dorsal attention, salience ventral attention, control and default. In each path analysis, the direct and indirect effect of CMR_GLC_ on cognition was tested via its effect on fPET_SD_ and local and global efficiency (see Figure 1B for illustration). We also tested for a direct effect of fPET_SD_ on cognition. Finally, we tested for a further mediating role of fPET_SD_ on cognition via network efficiency.

For each model, goodness of fit was assessed as adequate if Chi-square was greater than p = .05. Standardised regression weights were tested at p-FDR < .05 in each model.

## Data Availability Statement

All statistics supporting the results of this study are given in the Supplementary Information. The datasets used and/or analysed during the current study available from the corresponding author on reasonable request.

## Acknowledgements

Jamadar is supported by an Australian National Health and Medical Research Council (NHMRC) Fellowship (APP1174164).

## Author Contributions

SDJ and GFE conceived the project. HD and SDJ, CM and GFE designed and developed the manuscript. HD and EL analysed the data. HD wrote the manuscript, SDJ, GFE and CM reviewed/edited the manuscript. All authors read and approved the final manuscript. We thank Robert Di Paolo, Gerard Murray, M. Navyaan Siddiqui, Katharina Voigt, Richard McIntyre, Lauren Hudswell and the staff at Monash Biomedical Imaging for their contributions to data acquisition and image reconstruction.

## Footnotes

1 Here we use the term ‘stationary’ to refer to a measure that is taken across the entire scan under the assumption of a steady state of brain function or physiology. This is consistent with the use of the term in other neuroimaging contexts, where stationarity is defined as a static or unchanging phenomenon across the measurement period [32] and with the concept of weak sense stationarity [48].

