## Supplementary Information for "Brain glucodynamic variability is an essential feature of the metabolism-cognition relationship"

~ Supplementary Methods and Results ~

**1. Supplementary Methods** **2**
1.1 Ethical Considerations 2
1.2 Participants 2
1.3 Cognitive Tests 2
1.4 Data acquisition 2
1.5 MR & PET Data Preprocessing 3
1.6 Data Analysis 3

**2. Supplementary Results** **4**
2.1. Sample characteristics 4
2.2. Principal Component Analysis of Cognitive Measures 5
2.3 Age Group Differences in $fPET_{SD}$ 5
2.4 Means and standard deviation of measures used in path analyses 5
2.5 Path analysis model fit for models with direct path from CMRGLC to cognition 6
2.6 Path analysis statistics for the direct and indirect effects of $CMR_{GLC}$ and $fPET_{SD}$ on network efficiency and cognition 6
2.7 Data Table for region-level effects of age and head motion 8
2.8 Data table for region-wise age differences in metabolic rates of glucose 9

### 1. Supplementary Methods

##### *1.1 Ethical Considerations*

This study was conducted in accordance with Australian Code for the Responsible Conduct of Research (2007) and the Australian National Statement on Ethical Conduct in Human Research (2007). The Monash Health Principal Medical Physicist Administration approved the use of ionizing radiation based on the Australian Radiation Protection and Nuclear Safety Agency Code of Practice (2005). For participants older than 18 years, there is an annual radiation exposure limit of 5 mSv, higher than the effective dose of radiation in this study of 4.9 mSv. Participants provided informed consent to participate in the study.

##### *1.2 Participants*

Local advertising was used to recruit ninety participants from the general community. A screening interview was conducted to ensure that participants had the capacity to provide informed consent. Participants were also screened to ensure that they did not have a diagnosis of diabetes, neurological or psychiatric illness, claustrophobia or non-MR compatible implants. They were also excluded if they had received a clinical or research PET scan in the past 12 months. Women were screened for current or suspected pregnancy. Participants received a $100 voucher for participating in the study. Twelve participants were excluded from further analyses due to blood haemolysis or well counter issues preventing kinetic modelling (N=8), excessive head motion (N=2) or incomplete PET scan or image reconstruction (N=2).

##### *1.3 Cognitive Tests*

**Hopkins Verbal Learning Test (HVLT).** The HVLT is a three-trial list learning and free recall task. The learning trials comprised 12 words, four words from each of three semantic categories [1]. Approximately 20–25 minutes after the learning trails, participants completed delayed recall and recognition trials. The delayed recall required free recall of any of the 12 words. The recognition trial comprised 24 words, including the 12 target words and 12 false-positives, six semantically related, and six semantically unrelated. Delayed recall was calculated as the total words recalled.

**Task Switching.** For task switching, a computerised test was used in which participants were presented with a word and had to perform a categorisation task. The categorisation task was dependant on two cues that appeared on screen across the trials. One cue was a heart symbol, for which participants were asked to categorise the word presented via a key press as either a LIVING or a NON-LIVING object. If the cue was an arrow-cross, participants were asked to categorise the word as either BIGGER or SMALLER than a basketball. The cue was randomised across trials. Half the trials were switch trials and half were non-switch trials. Half the switch and non-switch trials was congruent in the key presses for either task, half was incongruent. The task switching measure was latency of correctly responding to a switch trial [2].

**Stop Signal.** The stop signal trial was a computer-based test [3]. Participants were required to press the left response key if an arrow on screen pointed left and the right response key if the arrow pointed right. If a signal beep sounded, participants were instructed stop their response. The delay between presentation of an arrow and signal beep started at 250ms and was altered up or down by 50ms based on performance. The delay increased up to 1150ms if the previous stop signal trial was successful and decreased down to 50ms if the previous stop signal trial was unsuccessful. The stimulus onset asynchrony between the onset of a fixation circles at the start of each trial was 2000ms. Reaction time in the stop signal trials was recorded.

**Digit Symbol Substitution.** A computer-based task presenting participants with a matric of 18 column and 16 rows [4]. Participants were required to translate symbols shown above the matrix in a key into digits in the matrix. The trail lasted two minutes. Performance was measured as total count of correct responses.

##### *1.4 Data acquisition*

Participants underwent a 90-minute simultaneous MR-PET scan in a Siemens (Erlangen) Biograph 3-Tesla molecular MR scanner. Participants were instructed to consume a high-protein and low-sugar diet for 24 hours and to fast completely for six hours prior to the scan. They were also asked to consume 2–6 glasses of water in the six hours before the scan. Upon arrival, participants were cannulated in the vein in each forearm and a baseline blood sample of 10ml was taken. At the start of the scan, half of the 260 MBq FDG tracer was administered as a bolus to provide a strong PET signal. The remaining 130 MBq of the tracer dose was infused over 50 minutes at a rate of 36ml/hour. This combined bolus plus constant infusion protocol provides a good balance between a rapid increase in signal-to-noise ratio at the beginning of the scan, and maintenance of signal-to-noise ratio over the length of the scan [5].

Participants were positioned supine in the scanner bore and instructed to lie as still as possible. Their head was placed in a 32-channel radiofrequency head coil. The scan sequence commenced with non-functional T1 and T2 MRI scans in the first 12 minutes to image the anatomical grey and white matter structures, respectively. The T1 3DMPRAGE scan parameters were: TA = 3.49 min, TR = 1,640ms, TE = 234ms, flip angle = 8°, field of view = 256 × 256 $mm^2$, voxel size = 1.0 × 1.0 × 1.0 $mm^3$, 176 slices, sagittal acquisition. The T2 FLAIR parameters were: TA = 5.52 min, TR = 5,000ms, TE = 396ms, field of view = 250 × 250 $mm^2$, voxel size = .5 × .5 × 1 $mm^3$, 160 slices). Thirteen minutes into the scan, list-mode PET (voxel size = 1.39 x 1.39 x 5.0$mm^3$) and T2* EPI BOLD-fMRI (TA = 40 minutes; TR = 1,000ms, TE = 39ms, FOV = 210 mm2, 2.4 × 2.4 × 2.4 $mm^3$ voxels, 64 slices, ascending axial acquisition) sequences were initiated. A 40-minute resting-state scan was undertaken while participants watched a movie of a drone flying over the Hawaii

Islands. Pseudo-continuous arterial spin labelling (pc-ASL) and diffusion-weighted imaging (DWI) was also acquired at the end of the scan but are not reported here.

Participants' plasma radioactivity levels were measured across the scan. Beginning at 10-minutes post infusion, 5ml blood samples were taken at 10 minute intervals from the right forearm. The blood sample were directly placed in a centrifuge and spun for 5 minutes at 2,000 rpm (RCF ~ 515g). 1,000-µL plasma was pipetted to a counting tube and placed in a well counter for four minutes. For each sample, the count start time, total number of counts, and counts per minute were documented.

##### *1.5 MR & PET Data Preprocessing*

For the T1 images, the brain was extracted in Freesurfer and registered to MNI152 space. The quality of the pial and white matter surface was manually checked and corrected if needed.

Participants' the list-mode PET data for were binned into 344 3D sinogram frames at 16s intervals. Attenuation was corrected via the pseudo-CT method for hybrid PET-MR scanners [6]. 3D volumes were constructed from the sinogram frames using the Ordinary Poisson-Ordered Subset Expectation Maximization algorithm (3 iterations, 21 subsets) with point spread function correction. The reconstructed DICOM slices were converted to NIFTI format. The NIFTI files were 344 × 344 × 127 (size: 1.39 × 1.39 × 2.03 $mm^3$) for each volume. A single 4D NIFTI volume was formed by temporally concatenating the 3D volumes. The 4D PET volumes were motion corrected [7], using the mean PET image to mask the data. PET images were corrected for partial volume effects using the modified Müller-Gartner method [8]. A grey matter threshold of 20-30% is recommended in ageing due to atrophy in older adults [8]. We used a 25% grey matter threshold and surface-based spatial smoothing [9]. To increase the signal-to-noise ratio, a Gaussian kernel was used with a full width at half maximum of 8 mm.

The relative head displacement was extracted from participants' head motion correction reports as the mean head movement in each frame relative to the following frame. Framewise displacement values were calculated to index the effects of translational and rotational head displacement, or the amount of head movement between consecutive frames. Specifically, framewise displacement was calculated as the absolute sum of three translation and three rotation length derivates [10]. The fPET timeseries for participants' were denoised by regressing out white matter and CSF signals. A low pass filter (.0625 Hz) was used to filter out high frequency noise [11, 12]. Regions of interest were generated for the 100 regions of the Schaefer Atlas [13].

##### *1.6 Data Analysis*

**Standard deviation of the timeseries ($fPET_{SD}$).** $fPET_{SD}$ was calculated from the individual participant's 100 regional timeseries. Data from 20 to 50 minutes of the scan was used to reduce the impact of the initial FDG uptake ramp and decrease post cessation of the constant infusion [5]. $fPET_{SD}$ measures the dispersion or spread of the signal around the mean, reflecting variability across the timeseries (see Figure 1A for illustration).

**Graph metrics.** We used the global and local efficiency graph metrics calculated at the top 30% of network edges based on relative edge strength. These metrics and threshold were chosen because they have been shown to have validity for studying network and cognitive changes in ageing using fPET [14, 15] and fMRI [16]. Local efficiency at each node is the average of shortest inverse-distances between the node and its neighbouring node in the local sub-graph. Global efficiency at each node is the average of the shortest inverse-distances between the node and all other nodes in the entire graph. Local and global efficiency across the whole graph are measures of local and global integration, respectively [17]. Global and local efficiency were also calculated for the regions of the dorsal attention, salience ventral attention, control and default networks for the sub-graph of those networks. We focused our analyses on these networks given their central role in the higher order cognitive processes [18-22]; and on the basis of our previous results in this dataset showing that $CMR_{GLC}$ in primary processing networks has limited association with cognitive performance [23].

**Cerebral Metabolic Rates of Glucose.** Regional $CMR_{GLC}$ was indexed as a 'stationary' measure of glucose metabolism across the entire scan. Calculations of regional $CMR_{GLC}$ were undertaken using the FDG time activity curves for the Schaefer atlas regions [24]. Participant's baseline plasma glucose (mmol) and their decay-corrected plasma FDG values were entered as an input function to Patlak models. The FDG and plasma data were used from the 10 minute point onwards, corresponding to the point of stable signal and the period of the fPET timeseries noted above. The regional $CMR_{GLC}$ values for two younger and one older participant were found to be more than 3.0 standard deviations from the mean and were exclude from analyses in which $CMR_{GLC}$ was used.

**Principal Component Analysis of Cognitive Measures.** Principal components analysis was used to reduce dimensionality in the cognitive data. The four cognitive measures were converted to Z-scores and entered in a Principal Component Analysis (PCA) with varimax rotations for components with eigenvalues greater than one. Participant component scores were saved for use in further analyses.

**Statistical Analysis.** To test our hypothesis that older age is associated with lower variability of the fPET timeseries, a series of general linear models (GLMs) was run in which regional $fPET_{SD}$ was the dependent variable. Given that head motion may impact variability of the fPET timeseries, we included it as a covariate in the analyses. Partial eta squared ($\eta^2_p$) was used to quantify the effect sizes in the GLMs. Each main effect was FDR-corrected at $p < .05$ across the 100 regions.

We used path analyses to test our hypotheses that that lower $fPET_{SD}$ will be associated with lower network efficiency and that $fPET_{SD}$ mediates the associated between 'stationary' glucose metabolism, network efficiency and cognition. To control for head motion, head motion (framewise displacement) was regressed on $fPET_{SD}$ and the residuals saved for each participant.

A separate path analysis was undertaken at two spatial scales: 1) the whole brain; and 2) each of the four networks separately: dorsal attention, salience ventral attention, control, default. For the analyses at the network level, the average network $fPET_{SD}$, $CMR_{GLC}$ and local and global efficiency was calculated from the regional values. The regions within the networks differ in cortical volume. Hence, each regional value was weighted by the percentage its volume represented in the total network volume.

In each path analysis, the direct and indirect effect of $CMR_{GLC}$ on cognition was tested via its effect on $fPET_{SD}$ and local and global efficiency (see Figure 1B for illustration). We also tested for a direct effect of $fPET_{SD}$ on cognition. Finally, we tested for a further mediating role of $fPET_{SD}$ on cognition via network efficiency.

For each model, goodness of fit was assessed as adequate if Chi-square was greater than p = .05. Standardised regression weights were tested at p-FDR < .05 in each model.

#### 2. Supplementary Results

##### *2.1. Sample characteristics*

The characteristics of the whole sample (N =78), as well as the younger (N = 35) and older (N = 43) participants, are shown in Table 1. The mean age of the whole sample was 54.0 years (SD = 24.6). The proportion of women was 53%. The average years of education was 17. Average BMI was 25.0 kg/m$^2$, resting heart rate was 78 BPM and systolic and diastolic blood pressure were 135 and 82 mmHg, respectively. The mean fasting blood glucose was 5.0 mmol/L.

The mean age of the younger group was 27.8 years and the older group 75.5 years. The proportion of women in the younger group (60%) and older group (48%) was not significantly different. The average years of education was higher in the younger (18.0) than the older group (16.2). These years of education are slightly above the average adult population in Australia [25].

The older group had significantly higher mean systolic blood pressure than the younger group (148mmHg vs 119mmHg). The older group also had a higher fasting blood glucose level (5.2 vs 4.7 mmol/L). A total of 27 people in the older group and eight in the younger group met diagnostic guidelines for hypertension. These rates are on par with those in the Australian adult population [26]. Although older people had a slightly higher mean BMI than younger people (25.7 vs 24.3 kg/m$^2$), the difference was not statistically significant. Nine participants (12%) would meet the definition for obesity (BMI above 30), including three younger and six older participants. This is below the 32% of the Australian adult population that is obese [26]. Head motion was significantly higher in the older than the younger group and was used as a covariate in analyses of fPET signal variability (below).

Supplementary Table 1. *Demographics for the whole sample and comparison of older and younger groups. Continuous variables are mean (standard deviation); categorical variables are number and percentage.*

| | Whole Sample (N = 78) | | Younger (N = 35) | | Older (N = 43) | | Younger vs Older p-value[1] |
|---|---|---|---|---|---|---|---|
| | Mean | SD | Mean | SD | Mean | SD | |
| Age (years) | 54.0 | 24.6 | 27.8 | 6.4 | 75.4 | 5.7 | 0.000 |
| Sex (number and % female) | 41 (53%) | | 21 (60%) | | 20 (49%) | | 0.235 |
| Education (years) | 17.0 | 3.5 | 18.0 | 2.8 | 16.3 | 3.8 | 0.034 |
| Body Mass Index (kg/m2) | 25.0 | 4.1 | 24.3 | 4.7 | 25.7 | 3.5 | 0.139 |
| Systolic Blood Pressure (mmHg) | 135.2 | 25.7 | 119.1 | 18.1 | 148.3 | 23.5 | 0.000 |
| Diastolic Blood Pressure (mmHg) | 81.6 | 12.9 | 78.9 | 13.5 | 83.8 | 12.2 | 0.099 |
| Resting Heart Rate (bpm) | 77.9 | 15.8 | 82.4 | 17.6 | 74.3 | 13.2 | 0.024 |
| Fasting blood glucose (mmol/L) | 5.0 | 0.5 | 4.7 | 0.4 | 5.2 | 0.5 | 0.000 |
| HVLT: Delayed Recall | 8.1 | 2.8 | 9.3 | 2.5 | 7.2 | 2.6 | 0.000 |
| Category Switch: RT in Switch Trials | 1.8 | 0.7 | 1.4 | 0.4 | 2.2 | 0.7 | 0.000 |
| Stop Signal: RT in Stop Signal Trials | 0.57 | 0.12 | 0.53 | 0.12 | 0.60 | 0.12 | 0.007 |
| Digit Symbol Substitution: Correct Count | 44.7 | 23.0 | 65.3 | 13.9 | 28.4 | 13.9 | 0.000 |
| Head Motion (Framewise Displacement) | 0.3 | 0.1 | 0.2 | 0.1 | 0.3 | 0.1 | 0.017 |

[1]P-values are based on T-test for continuous (2-sided) and Ch-square for categorical variables. Note: Education was not available for two younger and one older adult. One older participant's digit symbol substitution score was more than 3 standard deviations below the mean, and their data was excluded from analyses of cognition.

##### 2.2. Principal Component Analysis of Cognitive Measures

The principal components analysis yielded one significant component with an eigenvalue greater than one, explaining 57.6% of the variance. The component loadings were: HVLT delayed recall, .800; category switch reaction time, .802; stop signal reaction time, .602 and digit symbol substitution count, .812.

##### 2.3 Age Group Differences in $fPET_{SD}$

The descriptive statistics and GLMs testing for age group differences are available in Supplement 2.7 and 2.8. Older age was associated with lower $fPET_{SD}$ in 74 regions. There was no region in which older adults had higher $fPET_{SD}$ than younger adults. The regions showing the largest age group differences were the lateral prefrontal cortex (effect sizes ranging .158 to .164), inferior parietal lobule (.143) and precuneus (.166) in the control network; the dorsal prefrontal (.195 to .243), medial prefrontal (.148) and ventral prefrontal (.142 to .169) cortices, as well as the parietal lobule (.172) in the default network; the insula (.141 to .190) and medial (.167) and lateral (.159) prefrontal cortices in the salience ventral attention network; the temporal pole (.168) and orbital prefrontal cortex (.147) in the limbic network; the superior parietal lobule (.132 to .188) and post central gyrus (.142) in the dorsal attention network; and several regions In the somatomotor network (.152 to .221). Head motion was also associated with $fPET_{SD}$ in 21 regions.

##### 2.4 Means and standard deviation of measures used in path analyses

Supplementary Table 2. *Young and older adult group means (and standard deviation) on the measures used in the path analyses and t-tests of group differences. The means are plotted in Figure 2 in the main manuscript.*

| | Younger Adults | | Older Adults | | T-test: Younger vs Older | | |
|---|---|---|---|---|---|---|---|
| | Mean | SD | Mean | SD | t-value | p-value | p-FDR |
| $CMR_{GLC}$ | | | | | | | |
| Whole Brain | 3.56 | 0.88 | 2.96 | 0.87 | 2.9 | 0.005 | NA |
| Dorsal Attention | 3.18 | 0.78 | 2.71 | 0.74 | 2.6 | 0.011 | 0.011 |
| Salience Ventral Attention | 3.65 | 0.89 | 2.88 | 0.88 | 3.8 | 0.000 | 0.000 |
| Control | 3.79 | 0.95 | 3.13 | 0.96 | 2.9 | 0.004 | 0.005 |
| Default | 3.82 | 0.95 | 3.06 | 0.92 | 3.5 | 0.001 | 0.002 |
| Residual $fPET_{SD}$ (Head Motion Regressed Out) | | | | | | | |
| Whole Brain | 24.7 | 47.5 | -20.6 | 59.4 | 3.6 | 0.000 | NA |
| Dorsal Attention | 18.8 | 40.3 | -19.3 | 53.4 | 3.5 | 0.001 | 0.001 |
| Salience Ventral Attention | 30.0 | 62.0 | -27.4 | 75.9 | 3.6 | 0.001 | 0.001 |
| Control | 24.3 | 57.1 | -28.0 | 65.9 | 3.7 | 0.000 | 0.000 |
| Default | 22.4 | 53.8 | -24.4 | 64.9 | 3.4 | 0.001 | 0.001 |
| Global Efficiency | | | | | | | |
| Whole Brain | 0.618 | 0.037 | 0.632 | 0.040 | -1.6 | 0.213 | NA |
| Dorsal Attention | 0.508 | 0.087 | 0.584 | 0.068 | -4.3 | 0.000 | 0.000 |
| Salience Ventral Attention | 0.555 | 0.049 | 0.584 | 0.065 | -2.2 | 0.031 | 0.062 |
| Control | 0.669 | 0.048 | 0.681 | 0.041 | -1.2 | 0.226 | 0.226 |
| Default | 0.599 | 0.065 | 0.625 | 0.057 | -1.9 | 0.066 | 0.088 |
| Local Efficiency | | | | | | | |
| Whole Brain | 0.747 | 0.052 | 0.712 | 0.050 | 3 | 0.004 | NA |
| Dorsal Attention | 0.626 | 0.108 | 0.666 | 0.085 | -1.8 | 0.074 | 0.074 |
| Salience Ventral Attention | 0.716 | 0.067 | 0.680 | 0.077 | 2.2 | 0.034 | 0.068 |
| Control | 0.783 | 0.040 | 0.753 | 0.043 | 3.2 | 0.002 | 0.008 |
| Default | 0.719 | 0.056 | 0.695 | 0.055 | 1.9 | 0.057 | 0.074 |

Note: FDR-correction was undertaken across networks for each measure.

##### *2.5 Path analysis model fit for models with direct path from $CMR_{GLC}$ to cognition*

*Supplementary Table 3. Path analysis model fit comparing models with a direct path to $CMR_{GLC}$ rather than an indirect path via local and global efficiency and $fPET_{SD}$.*

| | Model: $CMR_{GLC}$ Indirect to Cognition Via Network Efficiency and $fPET_{SD}$ (per Supplement 2.6) | | Model: $CMR_{GLC}$ Direct to Cognition Rather than Via Network Efficiency and $fPET_{SD}$ | | | |
|---|---|---|---|---|---|---|
| | Model Fit | | Model Fit | | Path $CMR_{GLC}$ to Cognition | |
| | Chi-Sq. | p-value | Chi-Sq. | p-value | Beta | p-value |
| Local Efficiency and Cognition | | | | | | |
| Whole Brain | 0.14 | 0.714 | 0.68 | 0.410 | 0.050 | 0.670 |
| Dorsal Attention | 0.11 | 0.742 | 4.77 | 0.029 | 0.048 | 0.694 |
| Salience Ventral Attention | 0.80 | 0.371 | 1.08 | 0.298 | 0.116 | 0.375 |
| Control | 0.00 | 0.997 | 0.11 | 0.739 | 0.000 | 0.998 |
| Default | 0.61 | 0.436 | 0.31 | 0.578 | 0.107 | 0.404 |
| Global Efficiency and Cognition | | | | | | |
| Whole Brain | 0.10 | 0.755 | 1.08 | 0.299 | 0.040 | 0.753 |
| Dorsal Attention | 0.04 | 0.837 | 3.39 | 0.066 | -0.034 | 0.770 |
| Salience ventral Attention | 1.00 | 0.318 | 1.11 | 0.294 | 0.126 | 0.280 |
| Control | 0.02 | 0.886 | 0.00 | 0.961 | 0.019 | 0.879 |
| Default | 0.41 | 0.523 | 0.41 | 0.520 | 0.085 | 0.501 |

##### *2.6 Path analysis statistics for the direct and indirect effects of $CMR_{GLC}$ and $fPET_{SD}$ on network efficiency and cognition*

*Supplementary Table 4. Path analyses of direct and indirect effects of $CMR_{GLC}$ and $fPET_{SD}$ on network efficiency and cognition. The significant direct or indirect effects on cognition are summarised in Figure 3 in the main manuscript.*

| | Model Fit | | $CMR_{GLC}$ -> $fPET_{SD}$ | | | $CMR_{GLC}$ -> Efficiency | | |
|---|---|---|---|---|---|---|---|---|
| | Chi-Sq. | p-value | Beta | p | p-FDR | Beta | p | p-FDR |
| | Local Efficiency | | | | | | | |
| Whole Brain | 0.14 | 0.714 | 0.48 | 0.000 | 0.000 | -0.11 | 0.400 | 0.400 |
| Dorsal Attention | 0.11 | 0.74 | 0.44 | 0.000 | 0.000 | -0.28 | 0.026 | 0.043 |
| Salience Ventral Attention | 0.80 | 0.37 | 0.58 | 0.000 | 0.000 | 0.15 | 0.293 | 0.293 |
| Control | 0.00 | 1.00 | 0.52 | 0.000 | 0.000 | 0.04 | 0.742 | 0.742 |
| Default | 0.61 | 0.44 | 0.52 | 0.000 | 0.000 | 0.08 | 0.573 | 0.573 |
| | Global Efficiency | | | | | | | |
| Whole Brain | 0.10 | 0.755 | 0.47 | 0.000 | 0.000 | 0.14 | 0.277 | 0.277 |
| Dorsa Attention | 0.04 | 0.84 | 0.44 | 0.000 | 0.000 | -0.23 | 0.065 | 0.081 |
| Salience Ventral Attention | 1.00 | 0.32 | 0.55 | 0.000 | 0.000 | 0.15 | 0.259 | 0.259 |
| Control | 0.02 | 0.89 | 0.52 | 0.000 | 0.000 | 0.01 | 0.961 | 0.961 |
| Default | 0.41 | 0.52 | 0.51 | 0.000 | 0.000 | -0.08 | 0.532 | 0.532 |

*Supplementary Table 4 cont.*

| | $fPET_{SD}$ -> Efficiency | | | $fPET_{SD}$ ->Cognition | | | Efficiency ->Cognition | | |
|---|---|---|---|---|---|---|---|---|---|
| | Beta | p | p-FDR | Beta | p | p-FDR | Beta | p | p-FDR |
| | Local Efficiency | | | | | | | | |
| Whole Brain | 0.33 | 0.009 | 0.015 | 0.26 | 0.013 | 0.016 | 0.36 | 0.000 | 0.000 |
| Dorsal Attention | -0.03 | 0.823 | 0.828 | 0.36 | 0.001 | 0.003 | -0.02 | 0.828 | 0.828 |
| Salience Ventral Attention | 0.16 | 0.235 | 0.293 | 0.30 | 0.006 | 0.015 | 0.22 | 0.042 | 0.070 |
| Control | 0.26 | 0.049 | 0.061 | 0.28 | 0.009 | 0.015 | 0.33 | 0.002 | 0.005 |
| Default | 0.15 | 0.265 | 0.442 | 0.31 | 0.004 | 0.010 | 0.09 | 0.413 | 0.516 |
| | Global Efficiency | | | | | | | | |
| Whole Brain | -0.31 | 0.014 | 0.018 | 0.30 | 0.005 | 0.013 | -0.26 | 0.014 | 0.018 |
| Dorsa Attention | -0.10 | 0.400 | 0.400 | 0.30 | 0.005 | 0.008 | -0.30 | 0.005 | 0.008 |
| Salience Ventral Attention | -0.30 | 0.025 | 0.031 | 0.23 | 0.018 | 0.030 | -0.47 | 0.000 | 0.000 |
| Control | -0.06 | 0.641 | 0.801 | 0.36 | 0.000 | 0.000 | -0.11 | 0.290 | 0.483 |
| Default | -0.17 | 0.201 | 0.251 | 0.28 | 0.010 | 0.025 | -0.23 | 0.036 | 0.060 |

#### 2.7 Data Table for region-level effects of age and head motion

| | Younger | | Older | | Age Group | | | | Head Motion (Framewise Displacement) | | | |
|---|---|---|---|---|---|---|---|---|---|---|---|---|
| | Mean | SD | Mean | SD | F | p | p-FDR | η2p | F | p | p-FDR | η2p |
| Visual Central: Extra Striate Cortex 1 L | 450.9 | 50.5 | 435.7 | 71.5 | 0.5 | 0.469 | 0.474 | 0.007 | 1.0 | 0.309 | 0.347 | 0.014 |
| Visual Central: Extra Striate Cortex 2 L | 400.6 | 118.5 | 358.7 | 77.3 | 2.6 | 0.109 | 0.127 | 0.034 | 0.3 | 0.565 | 0.608 | 0.004 |
| Visual Central: Striate Cortex 1 L | 574.7 | 75.8 | 508.3 | 105.4 | 7.1 | 0.010 | 0.015 | 0.087 | 1.5 | 0.231 | 0.278 | 0.019 |
| Visual Central: Extra Striate Cortex 3 L | 337.2 | 66.7 | 296.6 | 52.0 | 6.1 | 0.016 | 0.024 | 0.076 | 2.5 | 0.119 | 0.165 | 0.033 |
| Visual Peripheral: Extra Striate Inferior 1 L | 560.6 | 68.3 | 517.7 | 81.6 | 4.3 | 0.043 | 0.055 | 0.054 | 1.3 | 0.266 | 0.312 | 0.017 |
| Visual Peripheral: Striate Cortex Calcarine 1 L | 498.9 | 63.9 | 436.3 | 77.5 | 8.7 | 0.004 | 0.008 | 0.105 | 12.3 | 0.001 | 0.029 | 0.142 |
| Visual Peripheral: Extra Striate CortexSup 1 L | 460.9 | 78.1 | 399.3 | 66.3 | 9.7 | 0.003 | 0.007 | 0.116 | 3.8 | 0.055 | 0.104 | 0.049 |
| Somatomotor A: 1 L | 397.8 | 74.7 | 329.8 | 74.1 | 11.0 | 0.001 | 0.004 | 0.130 | 4.5 | 0.037 | 0.081 | 0.058 |
| Somatomotor A: 2 L | 369.4 | 63.8 | 303.7 | 60.1 | 15.9 | 0.000 | 0.001 | 0.177 | 3.9 | 0.052 | 0.100 | 0.050 |
| Somatomotor B: Auditory 1 L | 406.7 | 71.0 | 352.0 | 61.1 | 9.5 | 0.003 | 0.007 | 0.114 | 2.5 | 0.121 | 0.166 | 0.032 |
| Somatomotor B: S2 1 L | 799.0 | 119.5 | 646.5 | 135.5 | 20.2 | 0.000 | 0.000 | 0.215 | 4.5 | 0.038 | 0.081 | 0.057 |
| Somatomotor B: S2 2 L | 449.0 | 64.8 | 367.4 | 69.2 | 20.9 | 0.000 | 0.000 | 0.221 | 5.5 | 0.022 | 0.064 | 0.069 |
| Somatomotor B: Central 1 L | 494.1 | 73.3 | 423.5 | 84.8 | 9.0 | 0.004 | 0.008 | 0.108 | 12.1 | 0.001 | 0.029 | 0.140 |
| Dorsal Attention A: Temporal Occipital 1 L | 337.2 | 38.6 | 313.4 | 51.1 | 3.2 | 0.077 | 0.093 | 0.042 | 2.0 | 0.157 | 0.194 | 0.027 |
| Dorsal Attention A: Parietal Occipital 1 L | 438.7 | 66.4 | 407.9 | 64.5 | 1.9 | 0.170 | 0.185 | 0.025 | 5.2 | 0.026 | 0.069 | 0.066 |
| Dorsal Attention A: Superior Parietal Lobule 1 L | 365.0 | 48.9 | 314.2 | 60.2 | 11.3 | 0.001 | 0.003 | 0.132 | 4.0 | 0.048 | 0.095 | 0.052 |
| Dorsal Attention B: Post Central 1 L | 472.6 | 77.7 | 401.7 | 85.6 | 9.4 | 0.003 | 0.007 | 0.113 | 5.4 | 0.023 | 0.065 | 0.068 |
| Dorsal Attention B: Post Central 2 L | 608.3 | 117.6 | 509.6 | 117.7 | 8.8 | 0.004 | 0.008 | 0.106 | 5.6 | 0.021 | 0.064 | 0.070 |
| Dorsal Attention B: Post Central 3 L | 443.8 | 69.8 | 377.4 | 89.6 | 9.2 | 0.003 | 0.007 | 0.111 | 2.1 | 0.150 | 0.190 | 0.028 |
| Dorsal Attention B: Frontal Eye Fields 1 L | 432.4 | 73.6 | 360.7 | 80.9 | 12.1 | 0.001 | 0.002 | 0.141 | 2.3 | 0.138 | 0.179 | 0.030 |
| Salience Ventral Attention A: Parietal Operculum 1 L | 462.9 | 84.0 | 388.2 | 83.7 | 9.4 | 0.003 | 0.007 | 0.112 | 10.0 | 0.002 | 0.032 | 0.119 |
| Salience Ventral Attention A: Insula: 1 L | 466.9 | 60.0 | 394.4 | 71.5 | 17.3 | 0.000 | 0.001 | 0.190 | 2.7 | 0.107 | 0.152 | 0.035 |
| Salience Ventral Attention A: Insula: 2 L | 604.6 | 94.0 | 515.7 | 89.3 | 13.8 | 0.000 | 0.002 | 0.157 | 1.9 | 0.175 | 0.214 | 0.025 |
| Salience Ventral Attention A: Parietal Medial 1 L | 673.8 | 109.3 | 575.6 | 126.3 | 8.3 | 0.005 | 0.009 | 0.101 | 6.1 | 0.016 | 0.061 | 0.076 |
| Salience Ventral Attention A: Frontal Medial 1 L | 605.7 | 101.5 | 505.0 | 124.4 | 9.4 | 0.003 | 0.007 | 0.112 | 8.0 | 0.006 | 0.041 | 0.098 |
| Salience Ventral Attention B: Lateral Prefrontal Cortex 1 L | 456.8 | 61.2 | 378.8 | 84.5 | 14.0 | 0.000 | 0.002 | 0.159 | 9.5 | 0.003 | 0.037 | 0.113 |
| Salience Ventral Attention B: Medial Posterior Prefrontal 1 L | 474.8 | 67.3 | 408.9 | 89.8 | 8.0 | 0.006 | 0.010 | 0.097 | 7.2 | 0.009 | 0.046 | 0.088 |
| Limbic A Temporal Pole 1 L | 403.4 | 73.5 | 337.4 | 64.3 | 14.9 | 0.000 | 0.001 | 0.168 | 0.3 | 0.603 | 0.642 | 0.004 |
| Limbic A: Temporal Pole 2 L | 316.7 | 43.6 | 293.7 | 47.2 | 4.1 | 0.046 | 0.057 | 0.053 | 0.1 | 0.795 | 0.814 | 0.001 |
| Limbic B: Orbital Frontal Cortex 1 L | 389.9 | 48.8 | 370.6 | 64.5 | 1.7 | 0.192 | 0.203 | 0.023 | 0.1 | 0.797 | 0.814 | 0.001 |
| Control A: Intraparietal Sulcus 1 L | 430.5 | 70.7 | 354.2 | 73.8 | 17.4 | 0.000 | 0.001 | 0.190 | 0.7 | 0.402 | 0.446 | 0.010 |
| Control A: Lateral Prefrontal Cortex 1 L | 399.3 | 71.8 | 329.3 | 66.8 | 14.6 | 0.000 | 0.002 | 0.164 | 3.1 | 0.082 | 0.136 | 0.040 |
| Control A: Lateral Prefrontal Cortex 2 L | 490.8 | 68.7 | 412.7 | 79.4 | 14.3 | 0.000 | 0.002 | 0.162 | 8.1 | 0.006 | 0.041 | 0.099 |
| Control B: Lateral Prefrontal Cortexv 1 L | 423.7 | 74.2 | 370.9 | 69.2 | 6.3 | 0.015 | 0.022 | 0.078 | 6.2 | 0.015 | 0.061 | 0.077 |
| Control C: Precuneus 1 L | 696.5 | 97.5 | 612.6 | 108.4 | 8.9 | 0.004 | 0.008 | 0.107 | 2.5 | 0.116 | 0.163 | 0.033 |
| Control C: Precuneus 2 L | 613.9 | 95.5 | 510.3 | 102.3 | 14.8 | 0.000 | 0.001 | 0.166 | 5.6 | 0.021 | 0.064 | 0.070 |
| Control C: Cingulate Posterior 1 L | 675.9 | 111.1 | 601.4 | 124.4 | 4.4 | 0.040 | 0.054 | 0.056 | 4.7 | 0.034 | 0.080 | 0.060 |
| Default A: Dorsal Prefrontal Cortex 1 L | 632.9 | 116.2 | 516.5 | 120.4 | 12.6 | 0.001 | 0.002 | 0.145 | 6.5 | 0.013 | 0.057 | 0.080 |
| Default A: Precuneus Posterior Cingulate Cortex1 L | 568.9 | 102.7 | 516.3 | 108.4 | 1.9 | 0.175 | 0.187 | 0.025 | 8.4 | 0.005 | 0.041 | 0.102 |
| Default A: Medial Prefrontal Cortex 1 L | 431.3 | 56.4 | 388.0 | 70.0 | 5.2 | 0.025 | 0.035 | 0.066 | 4.7 | 0.033 | 0.080 | 0.060 |
| Default B: Temp 1 L | 332.6 | 48.9 | 306.9 | 50.8 | 5.1 | 0.028 | 0.038 | 0.064 | 0.1 | 0.705 | 0.742 | 0.002 |
| Default B: Temp 2 L | 468.1 | 58.9 | 432.0 | 67.3 | 3.0 | 0.090 | 0.106 | 0.038 | 7.2 | 0.009 | 0.046 | 0.089 |
| Default B: Inferior Parietal Lobule 1 L | 305.2 | 51.7 | 263.5 | 49.6 | 8.6 | 0.004 | 0.008 | 0.104 | 4.6 | 0.035 | 0.081 | 0.058 |
| Default B: Dorsal Prefrontal Cortex 1 L | 382.5 | 67.0 | 303.4 | 68.1 | 20.4 | 0.000 | 0.000 | 0.216 | 2.4 | 0.124 | 0.168 | 0.032 |
| Default B: Lateral Prefrontal Cortex 1 L | 528.2 | 85.1 | 462.4 | 96.9 | 5.6 | 0.020 | 0.029 | 0.071 | 7.3 | 0.009 | 0.046 | 0.089 |
| Default B: Ventral Prefrontal Cortex 1 L | 594.6 | 87.8 | 506.2 | 78.5 | 15.1 | 0.000 | 0.001 | 0.169 | 8.0 | 0.006 | 0.041 | 0.098 |
| Default B: Ventral Prefrontal Cortex 2 L | 381.7 | 72.2 | 319.4 | 65.0 | 12.3 | 0.001 | 0.002 | 0.143 | 1.3 | 0.259 | 0.309 | 0.017 |
| Default C: RetroSuperior Parietal Lobuleenial 1 L | 733.4 | 85.0 | 632.0 | 107.1 | 15.3 | 0.000 | 0.001 | 0.172 | 3.1 | 0.083 | 0.136 | 0.040 |
| Default C: Parahippocampal Cortex 1 L | 553.4 | 60.9 | 520.8 | 68.6 | 2.5 | 0.115 | 0.130 | 0.033 | 3.6 | 0.063 | 0.112 | 0.046 |
| Temporal Parietal 1 L | 496.7 | 79.4 | 435.3 | 73.5 | 9.5 | 0.003 | 0.007 | 0.114 | 1.1 | 0.299 | 0.340 | 0.015 |
| Visual Central: Extra Striate Cortex 1 R | 411.3 | 56.3 | 399.9 | 62.8 | 0.1 | 0.748 | 0.748 | 0.001 | 3.0 | 0.087 | 0.139 | 0.039 |
| Visual Central: Extra Striate Cortex 2 R | 381.7 | 99.0 | 348.1 | 73.2 | 1.4 | 0.239 | 0.249 | 0.019 | 2.8 | 0.097 | 0.144 | 0.037 |
| Visual Central: Extra Striate Cortex 3 R | 395.3 | 63.0 | 345.1 | 56.5 | 9.4 | 0.003 | 0.007 | 0.113 | 3.4 | 0.068 | 0.119 | 0.044 |
| Visual Peripheral: Striate Cortex Calcarine 1 R | 489.5 | 64.4 | 448.2 | 79.0 | 3.5 | 0.065 | 0.079 | 0.045 | 3.8 | 0.056 | 0.104 | 0.048 |
| Visual Peripheral: Extra Striate Inferior 1 R | 456.7 | 60.4 | 388.5 | 59.3 | 18.4 | 0.000 | 0.001 | 0.199 | 4.8 | 0.032 | 0.080 | 0.060 |
| Visual Peripheral: Extra Striate Superior 1 R | 385.9 | 59.1 | 339.3 | 63.4 | 7.4 | 0.008 | 0.013 | 0.091 | 3.0 | 0.088 | 0.139 | 0.039 |
| Somatomotor A: 1 R | 788.6 | 157.9 | 664.9 | 147.3 | 7.8 | 0.007 | 0.011 | 0.096 | 6.9 | 0.010 | 0.049 | 0.086 |
| Somatomotor A: 2 R | 489.1 | 72.9 | 408.8 | 84.2 | 13.9 | 0.000 | 0.002 | 0.158 | 4.9 | 0.030 | 0.077 | 0.062 |
| Somatomotor A: 3 R | 573.6 | 72.1 | 520.7 | 101.9 | 4.3 | 0.042 | 0.055 | 0.055 | 2.3 | 0.135 | 0.179 | 0.030 |
| Somatomotor A: 4 R | 383.9 | 64.5 | 336.1 | 81.1 | 4.7 | 0.033 | 0.045 | 0.060 | 4.6 | 0.036 | 0.081 | 0.058 |
| Somatomotor B: Auditory 1 R | 380.5 | 51.5 | 338.1 | 54.2 | 7.3 | 0.009 | 0.014 | 0.089 | 8.7 | 0.004 | 0.041 | 0.105 |
| Somatomotor B: S2 1 R | 683.5 | 106.1 | 572.1 | 113.2 | 13.3 | 0.000 | 0.002 | 0.152 | 7.8 | 0.007 | 0.042 | 0.095 |
| Somatomotor B: S2 2 R | 517.5 | 58.2 | 422.9 | 76.2 | 28.8 | 0.000 | 0.000 | 0.280 | 3.1 | 0.083 | 0.136 | 0.040 |
| Somatomotor B: Central 1 R | 520.3 | 63.8 | 469.0 | 103.9 | 3.1 | 0.082 | 0.098 | 0.040 | 8.0 | 0.006 | 0.041 | 0.097 |
| Dorsal Attention A: Temporal Occipital 1 R | 343.2 | 47.1 | 319.8 | 51.3 | 2.5 | 0.115 | 0.130 | 0.033 | 2.1 | 0.154 | 0.192 | 0.027 |
| Dorsal Attention A: Parietal Occipital 1 R | 350.5 | 42.5 | 331.4 | 56.0 | 1.3 | 0.253 | 0.261 | 0.018 | 2.7 | 0.104 | 0.150 | 0.035 |
| Dorsal Attention A: Superior Parietal Lobule 1 R | 343.5 | 40.5 | 287.3 | 56.1 | 17.1 | 0.000 | 0.001 | 0.188 | 9.1 | 0.003 | 0.038 | 0.110 |
| Dorsal Attention B: Post Central 1 R | 552.4 | 73.9 | 472.7 | 91.8 | 12.3 | 0.001 | 0.002 | 0.142 | 3.6 | 0.062 | 0.112 | 0.046 |
| Dorsal Attention B: Post Central 2 R | 430.6 | 69.6 | 364.4 | 90.6 | 9.0 | 0.004 | 0.008 | 0.109 | 2.2 | 0.141 | 0.181 | 0.029 |
| Dorsal Attention B: Frontal Eye Fields 1 R | 456.6 | 72.3 | 377.9 | 91.6 | 10.7 | 0.002 | 0.004 | 0.127 | 10.9 | 0.001 | 0.030 | 0.129 |
| Salience Ventral Attention A: Parietal Operculum 1 R | 464.1 | 62.6 | 392.4 | 75.5 | 14.0 | 0.000 | 0.002 | 0.159 | 6.1 | 0.016 | 0.061 | 0.076 |
| Salience Ventral Attention A: Insula: 1 R | 436.8 | 54.4 | 380.3 | 62.3 | 12.1 | 0.001 | 0.002 | 0.141 | 5.5 | 0.022 | 0.064 | 0.069 |
| Salience Ventral Attention A: Parietal Medial 1 R | 653.2 | 114.7 | 562.2 | 120.4 | 5.9 | 0.018 | 0.026 | 0.074 | 15.7 | 0.000 | 0.017 | 0.175 |
| Salience Ventral Attention A: Frontal Medial 1 R | 526.5 | 88.5 | 441.2 | 114.8 | 7.6 | 0.007 | 0.012 | 0.093 | 10.5 | 0.002 | 0.030 | 0.124 |
| Salience Ventral Attention B: Inferior Parietal Lobule 1 R | 451.1 | 72.2 | 403.6 | 80.3 | 4.2 | 0.043 | 0.055 | 0.054 | 4.5 | 0.038 | 0.081 | 0.057 |
| Salience Ventral Attention B: Lateral Prefrontal Cortex 1 R | 478.1 | 83.6 | 409.5 | 86.4 | 8.6 | 0.004 | 0.008 | 0.105 | 2.9 | 0.091 | 0.140 | 0.038 |
| Salience Ventral Attention B: Medial Posterior Prefrontal 1 R | 552.5 | 96.3 | 445.2 | 103.3 | 14.9 | 0.000 | 0.001 | 0.167 | 10.5 | 0.002 | 0.030 | 0.124 |
| Limbic B: Orbital Frontal Cortex 0 R | 415.2 | 71.3 | 354.2 | 70.9 | 12.7 | 0.001 | 0.002 | 0.147 | 0.0 | 1.000 | 1.000 | 0.000 |
| Limbic A: Temporal Pole 1 R | 285.3 | 41.2 | 273.4 | 50.7 | 0.8 | 0.382 | 0.390 | 0.010 | 0.4 | 0.512 | 0.563 | 0.006 |
| Control A: Intraparietal Sulcus 1 R | 414.5 | 82.3 | 362.6 | 88.5 | 5.5 | 0.021 | 0.030 | 0.069 | 0.4 | 0.527 | 0.573 | 0.005 |
| Control A: Lateral Prefrontal Cortex 1 R | 447.8 | 72.1 | 386.8 | 85.5 | 6.6 | 0.012 | 0.019 | 0.082 | 7.5 | 0.008 | 0.045 | 0.092 |
| Control A: Lateral Prefrontal Cortex 2 R | 437.9 | 71.7 | 359.9 | 78.2 | 14.8 | 0.000 | 0.001 | 0.167 | 4.3 | 0.041 | 0.085 | 0.055 |
| Control B: Temporal 1 R | 360.9 | 53.2 | 325.7 | 58.7 | 6.4 | 0.014 | 0.021 | 0.079 | 0.1 | 0.766 | 0.798 | 0.001 |
| Control B: inferior parietal lobule 1 R | 440.5 | 80.2 | 365.8 | 74.3 | 12.4 | 0.001 | 0.002 | 0.143 | 5.7 | 0.019 | 0.064 | 0.072 |
| Control B: Lateral Prefrontal Cortexd 1 R | 492.0 | 83.5 | 401.3 | 94.4 | 13.6 | 0.000 | 0.002 | 0.155 | 6.5 | 0.013 | 0.057 | 0.081 |
| Control B: Lateral Prefrontal Cortexv 1 R | 391.7 | 75.1 | 322.8 | 68.4 | 13.9 | 0.000 | 0.002 | 0.158 | 1.2 | 0.269 | 0.312 | 0.017 |
| Control C: Cingulate Posterior 1 R | 699.4 | 100.7 | 607.2 | 116.3 | 8.8 | 0.004 | 0.008 | 0.106 | 5.7 | 0.020 | 0.064 | 0.071 |
| Control C: Precuneus 1 R | 492.3 | 71.1 | 444.5 | 87.6 | 4.1 | 0.046 | 0.057 | 0.053 | 3.0 | 0.089 | 0.139 | 0.039 |
| Default A: Inferior Parietal Lobule 1 R | 433.0 | 76.0 | 392.7 | 100.3 | 1.9 | 0.168 | 0.185 | 0.026 | 3.3 | 0.073 | 0.126 | 0.043 |
| Default A: Dorsal Prefrontal Cortex 1 R | 530.0 | 92.1 | 423.6 | 103.7 | 17.9 | 0.000 | 0.001 | 0.195 | 1.1 | 0.297 | 0.340 | 0.015 |
| Default A: Precuneus Posterior Cingulate Cortex 1 R | 677.6 | 118.2 | 613.6 | 135.0 | 2.5 | 0.120 | 0.135 | 0.032 | 4.2 | 0.044 | 0.091 | 0.054 |
| Default A: Medial Prefrontal Cortex 1 R | 451.8 | 70.8 | 378.0 | 78.1 | 12.8 | 0.001 | 0.002 | 0.148 | 5.8 | 0.019 | 0.064 | 0.072 |
| Default B: Dorsal Prefrontal Cortex 1 R | 415.5 | 66.1 | 326.8 | 72.0 | 23.7 | 0.000 | 0.000 | 0.243 | 5.2 | 0.025 | 0.069 | 0.066 |
| Default B: Ventral Prefrontal Cortex 1 R | 516.4 | 70.9 | 470.2 | 82.9 | 3.9 | 0.052 | 0.064 | 0.050 | 4.0 | 0.049 | 0.095 | 0.052 |
| Default B: Ventral Prefrontal Cortex 2 R | 487.6 | 79.7 | 407.9 | 89.6 | 12.2 | 0.001 | 0.002 | 0.142 | 2.8 | 0.098 | 0.144 | 0.037 |
| Default C: RetroSuperior Parietal Lobuleenial 1 R | 767.1 | 82.9 | 681.6 | 114.4 | 8.9 | 0.004 | 0.008 | 0.107 | 5.5 | 0.021 | 0.064 | 0.069 |
| Default C: Parahippocampal Cortex 1 R | 446.6 | 58.3 | 425.7 | 63.3 | 2.0 | 0.158 | 0.176 | 0.027 | 0.0 | 0.990 | 1.000 | 0.000 |
| Temporal Parietal 1 R | 388.8 | 31.9 | 359.1 | 49.8 | 5.6 | 0.020 | 0.029 | 0.071 | 5.1 | 0.027 | 0.072 | 0.064 |
| Temporal Parietal 2 R | 421.4 | 55.2 | 395.3 | 66.7 | 1.9 | 0.176 | 0.187 | 0.025 | 2.3 | 0.138 | 0.179 | 0.030 |
| Temporal Parietal 3 R | 473.5 | 64.0 | 417.8 | 63.9 | 10.4 | 0.002 | 0.005 | 0.123 | 2.8 | 0.097 | 0.144 | 0.037 |

#### 2.8 Data table for region-wise age differences in metabolic rates of glucose

| | Younger Adults (N = 35) | | Older Adults (N = 42) | | Age Group | | | |
|---|---|---|---|---|---|---|---|---|
| | Mean | SD | Mean | SD | F | p | p-FDR | η2p |
| Visual Central: Extra Striate Cortex 1 L | 3.7 | 1.0 | 3.4 | 1.0 | 1.3 | 0.254 | 0.267 | 0.018 |
| Visual Central: Extra Striate Cortex 2 L | 3.6 | 1.0 | 3.3 | 1.1 | 1.0 | 0.332 | 0.336 | 0.013 |
| Visual Central: Striate Cortex 1 L | 3.9 | 1.2 | 3.6 | 1.2 | 1.3 | 0.266 | 0.277 | 0.017 |
| Visual Central: Extra Striate Cortex 3 L | 3.2 | 0.8 | 2.9 | 0.8 | 3.1 | 0.085 | 0.099 | 0.040 |
| Visual Peripheral: Extra Striate Inferior 1 L | 3.8 | 1.0 | 3.3 | 1.0 | 4.2 | 0.043 | 0.054 | 0.055 |
| Visual Peripheral: Striate Cortex Calcarine 1 L | 3.5 | 1.0 | 3.1 | 0.9 | 3.7 | 0.060 | 0.072 | 0.048 |
| Visual Peripheral: Extra Striate CortexSup 1 L | 3.5 | 0.9 | 3.0 | 0.9 | 5.3 | 0.025 | 0.037 | 0.067 |
| Somatomotor A: 1 L | 3.2 | 0.8 | 2.7 | 0.7 | 8.8 | 0.004 | 0.009 | 0.108 |
| Somatomotor A: 2 L | 3.1 | 0.8 | 2.6 | 0.8 | 6.4 | 0.014 | 0.023 | 0.080 |
| Somatomotor B: Auditory 1 L | 3.6 | 0.9 | 3.1 | 0.9 | 6.7 | 0.012 | 0.022 | 0.084 |
| Somatomotor B: S2 1 L | 4.0 | 1.0 | 3.4 | 0.9 | 7.4 | 0.008 | 0.016 | 0.093 |
| Somatomotor B: S2 2 L | 3.7 | 0.9 | 2.9 | 1.0 | 14.5 | 0.000 | 0.002 | 0.165 |
| Somatomotor B: Central 1 L | 3.4 | 0.8 | 3.0 | 0.8 | 4.4 | 0.039 | 0.051 | 0.057 |
| Dorsal Attention A: Temporal Occipital 1 L | 3.4 | 0.9 | 3.0 | 0.9 | 4.4 | 0.040 | 0.052 | 0.057 |
| Dorsal Attention A: Parietal Occipital 1 L | 3.2 | 0.8 | 2.8 | 0.8 | 4.7 | 0.034 | 0.046 | 0.060 |
| Dorsal Attention A: Superior Parietal Lobule 1 L | 3.3 | 0.9 | 2.8 | 0.8 | 5.1 | 0.028 | 0.039 | 0.065 |
| Dorsal Attention B: Post Central 1 L | 3.0 | 0.9 | 2.5 | 0.7 | 9.6 | 0.003 | 0.007 | 0.117 |
| Dorsal Attention B: Post Central 2 L | 2.8 | 0.8 | 2.3 | 0.6 | 9.9 | 0.002 | 0.006 | 0.119 |
| Dorsal Attention B: Post Central 3 L | 2.7 | 0.7 | 2.3 | 0.6 | 8.0 | 0.006 | 0.013 | 0.099 |
| Dorsal Attention B: Frontal Eye Fields 1 L | 3.4 | 0.8 | 2.7 | 0.7 | 14.9 | 0.000 | 0.002 | 0.170 |
| Salience Ventral Attention A: Parietal Operculum 1 L | 3.6 | 0.9 | 2.9 | 0.9 | 9.8 | 0.003 | 0.006 | 0.118 |
| Salience Ventral Attention A: Insula: 1 L | 2.9 | 0.7 | 2.3 | 0.7 | 16.9 | 0.000 | 0.001 | 0.188 |
| Salience Ventral Attention A: Insula: 2 L | 4.5 | 1.1 | 3.5 | 1.1 | 15.0 | 0.000 | 0.002 | 0.170 |
| Salience Ventral Attention A: Parietal Medial 1 L | 3.5 | 0.8 | 2.9 | 0.9 | 7.6 | 0.007 | 0.015 | 0.094 |
| Salience Ventral Attention A: Frontal Medial 1 L | 3.8 | 0.9 | 3.0 | 0.9 | 14.0 | 0.000 | 0.002 | 0.161 |
| Salience Ventral Attention B: Lateral Prefrontal Cortex 1 L | 4.1 | 1.1 | 3.1 | 0.9 | 16.8 | 0.000 | 0.001 | 0.188 |
| Salience Ventral Attention B: Medial Posterior Prefrontal 1 L | 3.3 | 0.8 | 2.6 | 0.8 | 16.0 | 0.000 | 0.001 | 0.180 |
| Limbic A Temporal Pole 1 L | 4.2 | 1.1 | 3.4 | 1.0 | 10.3 | 0.002 | 0.006 | 0.124 |
| Limbic A: Temporal Pole 2 L | 2.6 | 0.6 | 2.3 | 0.7 | 6.4 | 0.014 | 0.023 | 0.080 |
| Limbic B: Orbital Frontal Cortex 1 L | 3.7 | 0.9 | 3.2 | 1.0 | 5.2 | 0.025 | 0.037 | 0.067 |
| Control A: Intraparietal Sulcus 1 L | 3.5 | 0.9 | 2.8 | 0.8 | 10.7 | 0.002 | 0.005 | 0.127 |
| Control A: Lateral Prefrontal Cortex 1 L | 3.8 | 1.0 | 3.1 | 0.9 | 11.2 | 0.001 | 0.004 | 0.133 |
| Control A: Lateral Prefrontal Cortex 2 L | 3.8 | 1.0 | 3.0 | 1.0 | 12.2 | 0.001 | 0.004 | 0.143 |
| Control B: Lateral Prefrontal Cortexv 1 L | 4.4 | 1.1 | 3.6 | 1.0 | 11.2 | 0.001 | 0.004 | 0.133 |
| Control C: Precuneus 1 L | 4.3 | 1.1 | 3.9 | 1.2 | 2.4 | 0.123 | 0.135 | 0.032 |
| Control C: Precuneus 2 L | 3.5 | 0.8 | 3.1 | 0.9 | 4.7 | 0.034 | 0.046 | 0.060 |
| Control C: Cingulate Posterior 1 L | 3.3 | 0.9 | 2.9 | 0.9 | 3.3 | 0.073 | 0.087 | 0.043 |
| Default A: Dorsal Prefrontal Cortex 1 L | 4.0 | 1.1 | 3.0 | 0.9 | 17.8 | 0.000 | 0.001 | 0.196 |
| Default A: Precuneus Posterior Cingulate Cortex1 L | 4.3 | 1.1 | 3.7 | 1.1 | 4.3 | 0.041 | 0.052 | 0.056 |
| Default A: Medial Prefrontal Cortex 1 L | 3.8 | 0.9 | 3.0 | 0.9 | 16.4 | 0.000 | 0.001 | 0.183 |
| Default B: Temp 1 L | 3.3 | 0.8 | 2.7 | 0.8 | 10.6 | 0.002 | 0.005 | 0.127 |
| Default B: Temp 2 L | 3.6 | 0.9 | 3.0 | 0.9 | 10.2 | 0.002 | 0.006 | 0.123 |
| Default B: Inferior Parietal Lobule 1 L | 3.5 | 0.9 | 2.9 | 1.0 | 6.9 | 0.010 | 0.020 | 0.086 |
| Default B: Dorsal Prefrontal Cortex 1 L | 4.0 | 1.0 | 3.0 | 0.9 | 24.2 | 0.000 | 0.001 | 0.249 |
| Default B: Lateral Prefrontal Cortex 1 L | 4.0 | 1.0 | 3.2 | 1.0 | 12.4 | 0.001 | 0.004 | 0.145 |
| Default B: Ventral Prefrontal Cortex 1 L | 4.0 | 1.1 | 3.1 | 0.9 | 16.9 | 0.000 | 0.001 | 0.188 |
| Default B: Ventral Prefrontal Cortex 2 L | 4.3 | 1.1 | 3.4 | 1.0 | 12.3 | 0.001 | 0.004 | 0.144 |
| Default C: RetroSuperior Parietal Lobuleenial 1 L | 4.1 | 1.1 | 3.5 | 1.2 | 4.0 | 0.049 | 0.061 | 0.052 |
| Default C: Parahippocampal Cortex 1 L | 3.0 | 0.7 | 2.5 | 0.9 | 6.0 | 0.017 | 0.027 | 0.075 |
| Temporal Parietal 1 L | 3.7 | 0.9 | 3.1 | 0.9 | 7.5 | 0.008 | 0.016 | 0.093 |
| Visual Central: Extra Striate Cortex 1 R | 3.7 | 1.0 | 3.4 | 1.0 | 1.1 | 0.291 | 0.300 | 0.015 |
| Visual Central: Extra Striate Cortex 2 R | 3.7 | 1.0 | 3.5 | 1.2 | 0.6 | 0.456 | 0.456 | 0.008 |
| Visual Central: Extra Striate Cortex 3 R | 3.2 | 0.8 | 3.0 | 0.9 | 1.8 | 0.180 | 0.192 | 0.024 |
| Visual Peripheral: Striate Cortex Calcarine 1 R | 4.1 | 1.2 | 3.8 | 1.3 | 1.1 | 0.298 | 0.304 | 0.015 |
| Visual Peripheral: Extra Striate Inferior 1 R | 3.3 | 0.9 | 2.9 | 0.9 | 5.0 | 0.028 | 0.039 | 0.065 |
| Visual Peripheral: Extra Striate Superior 1 R | 3.2 | 0.9 | 2.9 | 0.9 | 2.8 | 0.101 | 0.114 | 0.036 |
| Somatomotor A: 1 R | 3.9 | 0.9 | 3.2 | 0.9 | 8.7 | 0.004 | 0.010 | 0.106 |
| Somatomotor A: 2 R | 3.3 | 0.8 | 2.8 | 0.8 | 6.2 | 0.015 | 0.024 | 0.079 |
| Somatomotor A: 3 R | 2.7 | 0.6 | 2.4 | 0.7 | 5.1 | 0.027 | 0.038 | 0.066 |
| Somatomotor A: 4 R | 3.1 | 0.8 | 2.7 | 0.8 | 6.9 | 0.011 | 0.020 | 0.086 |
| Somatomotor B: Auditory 1 R | 3.4 | 0.8 | 2.9 | 0.9 | 5.4 | 0.023 | 0.035 | 0.069 |
| Somatomotor B: S2 1 R | 3.9 | 1.0 | 3.2 | 1.1 | 8.7 | 0.004 | 0.010 | 0.106 |
| Somatomotor B: S2 2 R | 3.6 | 0.9 | 2.8 | 1.1 | 11.2 | 0.001 | 0.004 | 0.133 |
| Somatomotor B: Central 1 R | 3.2 | 0.7 | 2.9 | 0.9 | 2.3 | 0.132 | 0.142 | 0.031 |
| Dorsal Attention A: Temporal Occipital 1 R | 3.3 | 0.8 | 2.9 | 0.9 | 2.7 | 0.106 | 0.117 | 0.035 |
| Dorsal Attention A: Parietal Occipital 1 R | 3.4 | 0.8 | 2.9 | 0.9 | 4.3 | 0.041 | 0.052 | 0.056 |
| Dorsal Attention A: Superior Parietal Lobule 1 R | 2.9 | 0.7 | 2.5 | 0.6 | 6.0 | 0.017 | 0.027 | 0.075 |
| Dorsal Attention B: Post Central 1 R | 3.1 | 0.9 | 2.6 | 0.8 | 6.5 | 0.013 | 0.023 | 0.082 |
| Dorsal Attention B: Post Central 2 R | 2.9 | 0.7 | 2.5 | 0.8 | 4.9 | 0.030 | 0.041 | 0.063 |
| Dorsal Attention B: Frontal Eye Fields 1 R | 3.3 | 0.8 | 2.6 | 0.7 | 14.8 | 0.000 | 0.002 | 0.169 |
| Salience Ventral Attention A: Parietal Operculum 1 R | 3.5 | 0.9 | 2.8 | 0.9 | 11.5 | 0.001 | 0.004 | 0.136 |
| Salience Ventral Attention A: Insula: 1 R | 3.7 | 0.9 | 3.0 | 1.0 | 10.1 | 0.002 | 0.006 | 0.121 |
| Salience Ventral Attention A: Parietal Medial 1 R | 3.6 | 0.9 | 3.1 | 1.0 | 6.4 | 0.014 | 0.023 | 0.080 |
| Salience Ventral Attention A: Frontal Medial 1 R | 3.7 | 0.9 | 2.9 | 0.9 | 13.6 | 0.000 | 0.002 | 0.157 |
| Salience Ventral Attention B: Inferior Parietal Lobule 1 R | 3.3 | 0.9 | 2.7 | 0.8 | 11.2 | 0.001 | 0.004 | 0.133 |
| Salience Ventral Attention B: Lateral Prefrontal Cortex 1 R | 4.0 | 1.0 | 3.2 | 1.0 | 13.3 | 0.000 | 0.003 | 0.154 |
| Salience Ventral Attention B: Medial Posterior Prefrontal 1 R | 3.5 | 0.9 | 2.5 | 0.9 | 22.3 | 0.000 | 0.001 | 0.234 |
| Limbic B: Orbital Frontal Cortex 0 R | 4.3 | 1.2 | 3.5 | 1.1 | 9.2 | 0.003 | 0.008 | 0.112 |
| Limbic A: Temporal Pole 1 R | 2.8 | 0.7 | 2.4 | 0.8 | 5.6 | 0.021 | 0.032 | 0.071 |
| Control A: Intraparietal Sulcus 1 R | 2.9 | 0.8 | 2.5 | 0.9 | 3.7 | 0.059 | 0.072 | 0.048 |
| Control A: Lateral Prefrontal Cortex 1 R | 3.8 | 1.0 | 3.1 | 0.9 | 11.8 | 0.001 | 0.004 | 0.139 |
| Control A: Lateral Prefrontal Cortex 2 R | 3.4 | 0.9 | 2.8 | 0.9 | 9.8 | 0.002 | 0.006 | 0.119 |
| Control B: Temporal 1 R | 3.6 | 0.9 | 3.0 | 1.0 | 6.4 | 0.013 | 0.023 | 0.081 |
| Control B: inferior parietal lobule 1 R | 3.7 | 0.9 | 3.1 | 1.1 | 7.3 | 0.009 | 0.017 | 0.091 |
| Control B: Lateral Prefrontal Cortexd 1 R | 4.1 | 1.0 | 3.3 | 1.1 | 12.3 | 0.001 | 0.004 | 0.144 |
| Control B: Lateral Prefrontal Cortexv 1 R | 4.1 | 1.1 | 3.4 | 1.0 | 9.6 | 0.003 | 0.007 | 0.116 |
| Control C: Cingulate Posterior 1 R | 3.7 | 1.0 | 3.2 | 1.1 | 5.0 | 0.028 | 0.039 | 0.064 |
| Control C: Precuneus 1 R | 4.1 | 1.1 | 3.7 | 1.2 | 3.0 | 0.089 | 0.102 | 0.039 |
| Default A: Inferior Parietal Lobule 1 R | 3.4 | 0.9 | 2.8 | 1.0 | 6.4 | 0.013 | 0.023 | 0.081 |
| Default A: Dorsal Prefrontal Cortex 1 R | 3.7 | 1.1 | 2.8 | 0.8 | 18.0 | 0.000 | 0.001 | 0.198 |
| Default A: Precuneus Posterior Cingulate Cortex 1 R | 4.5 | 1.1 | 4.0 | 1.3 | 3.0 | 0.086 | 0.099 | 0.040 |
| Default A: Medial Prefrontal Cortex 1 R | 3.9 | 0.9 | 3.0 | 0.9 | 16.8 | 0.000 | 0.001 | 0.187 |
| Default B: Dorsal Prefrontal Cortex 1 R | 3.9 | 1.0 | 3.0 | 0.9 | 20.3 | 0.000 | 0.001 | 0.218 |
| Default B: Ventral Prefrontal Cortex 1 R | 4.0 | 1.0 | 3.2 | 1.0 | 11.1 | 0.001 | 0.004 | 0.132 |
| Default B: Ventral Prefrontal Cortex 2 R | 4.2 | 1.1 | 3.3 | 1.1 | 13.9 | 0.000 | 0.002 | 0.160 |
| Default C: RetroSuperior Parietal Lobuleenial 1 R | 4.3 | 1.1 | 3.8 | 1.4 | 2.4 | 0.126 | 0.137 | 0.032 |
| Default C: Parahippocampal Cortex 1 R | 3.0 | 0.7 | 2.7 | 0.8 | 3.2 | 0.079 | 0.093 | 0.042 |
| Temporal Parietal 1 R | 2.7 | 0.6 | 2.2 | 0.7 | 10.1 | 0.002 | 0.006 | 0.121 |
| Temporal Parietal 2 R | 3.5 | 0.9 | 2.9 | 0.9 | 8.5 | 0.005 | 0.010 | 0.105 |
| Temporal Parietal 3 R | 3.2 | 0.8 | 2.7 | 0.9 | 6.0 | 0.017 | 0.027 | 0.075 |
